# Integrative Omics reveals changes in the cellular landscape of yeast without peroxisomes

**DOI:** 10.1101/2024.03.20.585854

**Authors:** Tjasa Kosir, Hirak Das, Marc Pilegaard Pedersen, Marco Anteghini, Silke Oeljeklaus, Vitor Martins dos Santos, Ida J. van der Klei, Bettina Warscheid

**Affiliations:** Molecular Cell Biology, Groningen Biomolecular Sciences and Biotechnology Institute (BBA) University of Groningen, PO Box 11103, 9300 CC Groningen, The Netherlands; Theodor Boveri-Institute, Biochemistry II, Faculty of Chemistry and Pharmacy, University of Würzburg, 97074 Würzburg, Germany; Laboratory of Systems and Synthetic Biology, Wageningen University & Research, Wageningen WE, The Netherlands; Lifeglimmer GmbH, Berlin, Germany; Department of Bioprocess Engineering, Wageningen University & Research, Wageningen WE, The Netherlands

**Author notes:** Corresponding authors: Bettina Warscheid,; ORCID 0000-0001-5096-1975; Ida J. van der Klei,; ORCID 0000-0001-7165-9679. These authors contributed equally to this work (Tjasa Kosir, Hirak Das).

**Keywords:** Peroxisome, yeast, proteome, transcriptome, *PEX* genes, β-oxidation, mitochondria

## Abstract

Peroxisomes are organelles that are crucial for cellular metabolism. However, these organelles play also important roles in non-metabolic processes, such as signalling. To uncover the consequences of peroxisome deficiency, we compared two extremes, namely *Saccharomyces cerevisiae* wild-type and *pex3* cells, which lack functional peroxisomes, employing transcriptomics and quantitative proteomics technology. Cells were grown on acetate, a carbon source that involves peroxisomal enzymes of the glyoxylate cycle and does not repress peroxisomal proteins. Transcripts of peroxisomal β-oxidation genes and the corresponding proteins were enhanced in *pex3* cells. Peroxisome-deficiency also caused reduced levels of membrane bound peroxins, while the soluble receptors Pex5 and Pex7 were enhanced at the protein level. In addition, we observed alterations in non-peroxisomal transcripts and proteins, especially mitochondrial proteins involved in respiration or import processes. Our results not only reveal the impact of the absence of peroxisomes in yeast, but also represent a rich resource of candidate genes/proteins that are relevant in peroxisome biology.

**Summary:** Omics comparison of wild-type and peroxisome-deficient (*pex3*) yeast cells uncovered processes that are affected by loss of peroxisomes. β-oxidation enzymes were upregulated, whereas most peroxins were decreased. Also, several non-peroxisomal transcripts/proteins were significantly altered. Our data represent a rich source of candidate genes connected to peroxisome biology.

## Introduction

Peroxisomes are organelles that occur in almost all eukaryotic cells. Their importance is illustrated by pathogenic mutations that can cause devastating peroxisomal diseases such as Zellweger spectrum disorders (Waterham et al., 2016).

On a cellular level, peroxisomes consist of a proteinaceous matrix, which contains enzymes involved in various metabolic pathways, and is bound by a single membrane. Common peroxisomal enzymes are hydrogen peroxide-producing oxidases, catalase and the β-oxidation pathway members. Depending on the organism, developmental stage and cell type, peroxisomes can also harbour a broad spectrum of other metabolic pathways (Bartoszewska et al., 2011; Hu et al., 2012; Wanders and Waterham, 2006). The peroxisomal membrane contains a highly diverse set of proteins. Most extensively studied are membrane bound peroxins (encoded by *PEX* genes), which are required for the biogenesis of the organelle (Jansen et al., 2021). Examples of other peroxisomal membrane proteins (PMPs) include solute transporters and proteins involved in organelle fission and inheritance.

Peroxisomes also fulfil crucial non-metabolic functions. For instance, mammalian peroxisomes can serve as intracellular signalling platforms in innate immunity and inflammation (Islinger et al., 2018). In addition, they are implicated in ageing (Fransen et al., 2013) and in antiviral response (Ferreira et al., 2023). Hence, a detailed understanding of the molecular mechanisms involved in peroxisome biology is important for human health and disease.

Yeasts are simple model organisms that are extensively used to study peroxisomes. Since the molecular mechanisms of many processes in peroxisome biology are conserved, the knowledge obtained by yeast peroxisome research can be translated to higher eukaryotes, including humans. Yeast research substantially contributed to the identification and characterization of peroxins (Akşit and van der Klei, 2018; Erdmann et al., 1997; Mast and Aitchison, 2018). Mutant screens were performed to isolate strains defective in various aspects of peroxisome biology. The analysis of yeast libraries by high throughput fluorescence microscopy (FM) and proteomics studies of isolated peroxisomes or affinity-purified protein complexes of peroxisomal proteins, mostly peroxins, resulted in the discovery of many additional proteins and processes related to peroxisome biology (David et al., 2013; Grunau et al., 2009; Marelli et al., 2004; Mast et al., 2016; Yifrach et al., 2022). Peroxisomes cannot function in isolation but must communicate and cooperate with their environment to exchange metabolites and coordinate cellular responses (Silva et al., 2020). For instance, in yeast, β-oxidation of fatty acids is confined to peroxisomes and results in the formation of acetyl-CoA, which is transferred to mitochondria for further oxidation via the citric acid cycle. Recent studies indicate that physical contacts between both organelles stimulate acetyl-CoA transfer between both organelles (Shai et al., 2018), Moreover, the small GTPase Arf1 was demonstrated to localize to these contacts, where it coordinates acetyl-CoA transfer (Enkler et al., 2023).

Importantly, peroxisomes also form physical contacts with other cell organelles, but the function of most of them is relatively unexplored (Wu et al., 2023). In general, however, our knowledge of the processes involved in the communication and cooperation of peroxisomes with their environment is still in its infancy.

The aim of our current study is to uncover which cellular processes are influenced by peroxisomes in yeast. To this end, we compared two extremes, namely *Saccharomyces cerevisiae* wild-type (WT) and *pex3* cells, by transcriptomics and whole cell proteomics analysis. Yeast *pex3* cells lack functional peroxisomes but are fully viable and capable of growing on several carbon sources except oleic acid (Erdmann et al., 1989). For our studies, we have chosen acetate as the sole carbon source, whose metabolism involves the peroxisomal enzymes of the glyoxylate cycle (Kunze and Hartig, 2013). Notably, we refrained from growing yeast cells on glucose because it represses the expression of many peroxisomal proteins (Kayikci and Nielsen, 2015).

We show that peroxisome-deficiency elicits specific responses at the transcriptome and proteome level. In addition to a significant increase in the levels of most peroxisomal β-oxidation enzymes, virtually all membrane bound peroxins were strongly reduced, while the soluble receptor proteins Pex5 and Pex7 showed a higher abundance in *pex3* cells. Importantly, several non-peroxisomal proteins were also either up or downregulated in *pex3* cells. Especially, several mitochondrial proteins involved in protein import and refolding as well as in mitochondrial respiration were enhanced. Moreover, we identified several other non-peroxisomal transcripts and proteins, including several poorly characterized ones, which were changed in abundance and hence are very attractive candidates for further studies on their connection with peroxisome biology.

## Results

We first validated the use of acetate as an appropriate carbon source for our studies. Although several enzymes involved in acetate metabolism [e.g. enzymes of the glyoxylate cycle and carnitine acetyl-CoA transferase (Cat2)] were previously shown to be peroxisomal in oleic acid grown *S. cerevisiae* (reviewed in Kunze and Hartig, 2013; addition Nakatsukasa et al., 2015; Rosenthal et al., 2020; Schummer et al., 2020; Yifrach et al., 2016) the localization of most of them has not been reported for acetate grown cells. Moreover, we could not exclude an alternative localization of these enzymes in acetate grown cells. One example is malate synthase 1 (Mls1), which is peroxisomal in oleic acid grown *S. cerevisiae* cells, but cytosolic upon growth on acetate (Kunze et al., 2002; Yifrach et al., 2016).

We studied the localization of citrate synthase 2 (Cit2), malate dehydrogenase 3 (Mdh3) and Cat2. Cat2 has a dual localization in mitochondria and peroxisomes (Elgersma et al., 1995). The three proteins of interest were tagged at the N- or C-terminus with a fluorescence protein (FP). Cells were grown in batch cultures on acetate and analyzed by FM. As shown in **Figure 1a**, all three N-terminally tagged proteins showed a punctuate pattern, in line with a peroxisomal localization. This pattern was lost when the FP was fused to the C-terminus, which can be explained by the interference of the C-terminal tag with the peroxisomal targeting signal 1 (PTS1) that is present at the extreme C-terminus of these proteins (**Figure 1a**). Although a minor fraction of the Cat2-GFP fluorescence was present in the cytosol, Cat2-GFP was predominantly observed in mitochondria when C-terminally tagged, in line with the finding that Cat2 also contains a mitochondrial targeting signal at the N-terminus (van Roermund, 1999).

**Figure 1.**
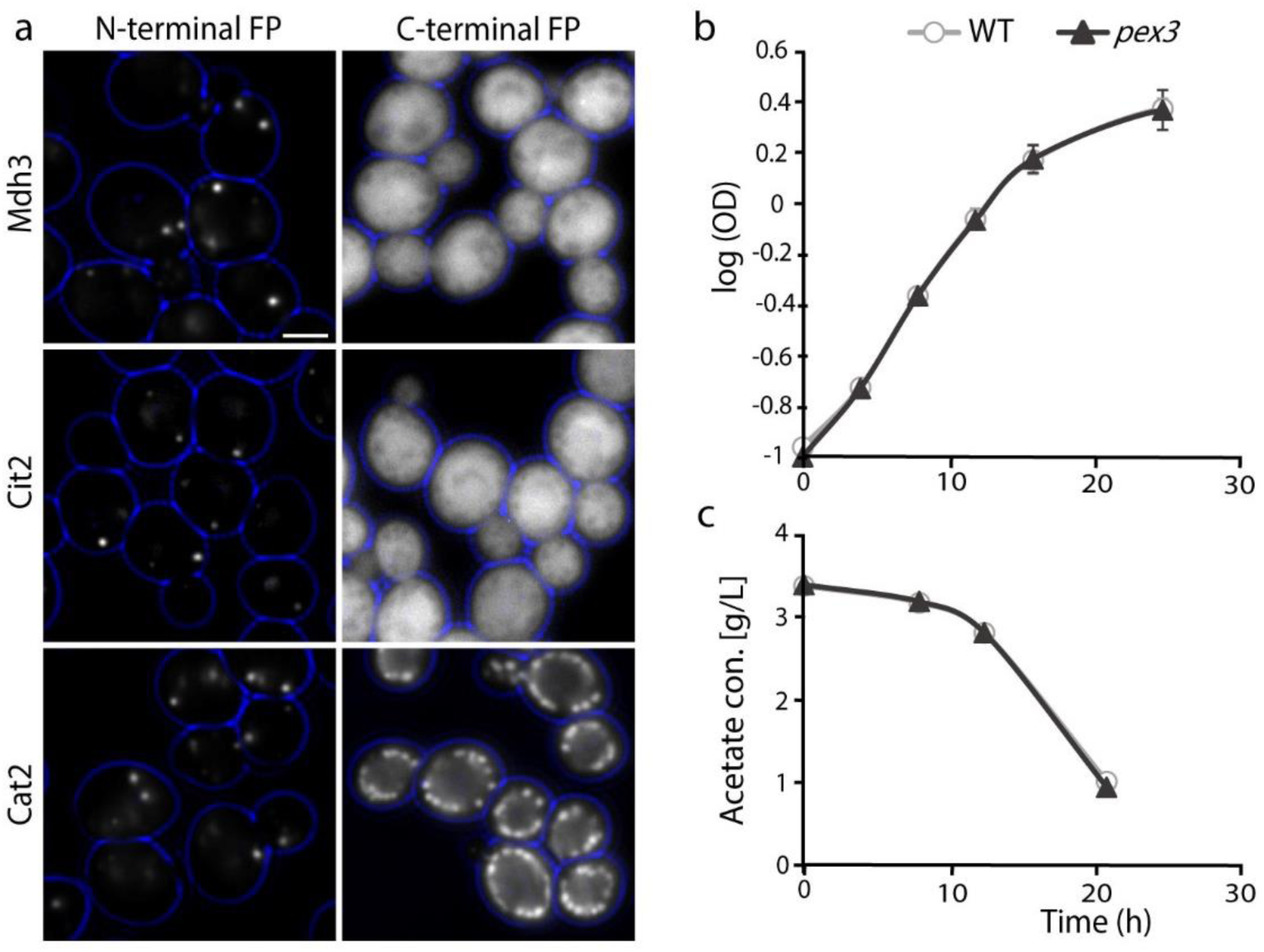
Validation of acetate as a suitable carbon source for the omics comparison of WT and *pex3* cells. (a) Fluorescence microscopy analysis of cells expressing Mdh3, Cit2 or Cat2 with the fluorescent protein (FP) GFP at the N-terminus under control of the *NOP1* promoter or tagged with the FP NeonGreen (Mdh3, Cat2) or GFP (Cit2) at the C-terminus under control of the endogenous promoter. Scale bar: 2.5 μm. (b) Growth curves of WT and *pex3* cells grown in batch cultures on acetate medium. The cell density is indicated as log optical density (OD) at 660 nm. The error bars represent the standard deviation (SD) from three independent experiments. (c) Acetate concentrations (g/L) in the media of WT and *pex3* cultures. The error bars represent SD from three independent experiments. Error bars are smaller than the symbols.

For a reliable comparison of the transcriptome and proteome of WT and *pex3* cells, it is essential that cell growth is identical to avoid artifacts caused by differences in growth parameters. As shown in **Figure 1b**, WT and *pex3* cells growth rates were similar in batch cultures containing acetate as the sole carbon source. Moreover, measurements of the residual concentrations of acetate in the growth medium revealed that acetate consumption was the same in both strains (**Figure 1c**).

To conclude, our data show that the loss of peroxisomes in *pex3* cells neither affects the growth of *S. cerevisiae* cells on acetate nor acetate consumption. Moreover, under these growth conditions, peroxisomes of WT cells harbor several enzymes involved in acetate metabolism and therefore the organelles are involved in the primary metabolism of the carbon source. This makes acetate a suitable carbon source for comparative quantitative omics studies of WT and *pex3* cells.

### Quantitative proteome and transcriptome analysis of WT *versus pex3* cells

To globally study the loss of peroxisomes in *S. cerevisiae* cells for its effect on the cellular proteome and transcriptome, WT and *pex3* cells were grown in batch cultures using acetate as a carbon source and harvested in the early exponential growth phase. Whole cell proteome and transcriptome analyses were performed from the identical samples, followed by quantitative data analysis and integrative evaluation of the obtained omics data (**Figure 2a**).

**Figure 2.**
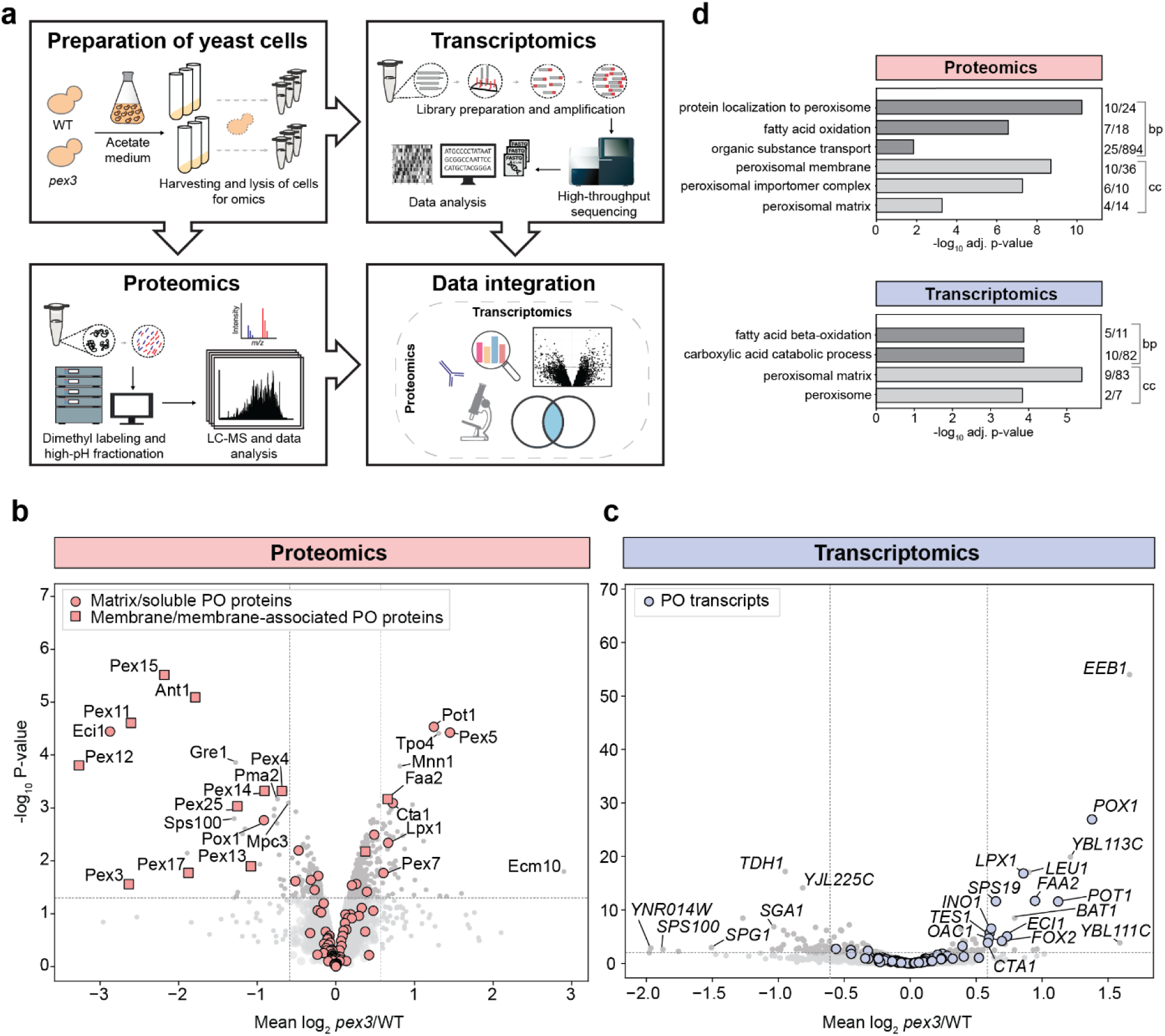
Proteomics and transcriptomics analysis of *S. cerevisiae* WT and *pex3* cells. (a) Outline of the experimental strategy employed for the systematic analysis of global changes in the proteome and transcriptome caused by the absence of functional peroxisomes in *pex3* cells. Experiments were performed in three independent biological replicates, and the same samples were analyzed by quantitative proteomics and transcriptomics. (b, c) Global differences in protein (b) and transcript (c) levels in *pex3 versus* WT cells. Mean log2 *pex3*/WT ratios were plotted against the corresponding log-transformed p-value (-log10) determined using the ‘linear models for microarray data’ (limma) approach (two-sided t-test; n=3) for the proteomics and the Wald test for the transcriptomics data. Horizontal and vertical lines indicate a p-value of 0.05 and a fold-change of ≤ 0.66 or ≥ 1.5, respectively. Information about a peroxisomal localization of proteins or association with peroxisomes is based on (David et al., 2022; Elbaz-Alon et al., 2014; Marelli et al., 2004; Motley et al., 2012; Tanaka et al., 2014; Yifrach et al., 2022). PO, peroxisomal and peroxisome-associated proteins or transcripts. (d) GO term overrepresentation analysis of proteins and transcripts with a fold-change of ≤ 0.66 or ≥ 1.5 (p-value < 0.05) for the domains “biological process” (bp) and “cellular component” (cc). Listed are selected terms with an adjusted (adj.) p-value of < 0.05. Numbers indicate the number of proteins or transcripts assigned to a given term and the number of proteins/transcripts with this term in the proteomics or transcriptomics dataset.

For quantitative whole cell proteomics of WT and *pex3* cells, we employed peptide stable isotope dimethyl labeling (Boersema et al., 2009), orthogonal peptide separation by high and low pH reversed-phase liquid chromatography (LC) followed by high-resolution mass spectrometry (MS). This in-depth analysis resulted in the quantification of 3,921 protein groups in at least two out of three replicates, of which 3,700 (94.4%) were quantified in all three replicates (**Table S1a**). We found 79 proteins with an average change in abundance of at least 1.5-fold (34/45 proteins reduced/increased; p-value of < 0.05, n=3) (**Figure 2b, Table S1a**). Using this proteomics approach, we obtained quantitative data for 92 peroxisomal proteins, with 17/11 reduced/increased in abundance (p-value < 0.05). Furthermore, most PMPs (10 out of 12, p-value < 0.05) were more than 1.5-fold reduced in abundance, whereas a few soluble peroxisomal proteins (5 out of 16, p-value < 0.05) were more than 1.5-fold increased in *pex3* cells (**Figure 2b** and **Table S1a**). Quantitative Western blotting of selected regulated proteins confirmed abundance changes in *pex3* cells determined by quantitative MS analysis as shown for the soluble peroxisomal protein Pot1 and the PTS1 receptor protein Pex5, which were both increased in *pex3* cells, and the PMP Pex14, which was 1.9-fold decreased. In addition, a 2.5-fold increase in abundance of the plasma membrane and vacuole transporter Tpo4 in *pex3* cells was confirmed by Western blotting (**Figures S1a** and **S1b**).

The complementary analysis of the transcriptome of WT and *pex3* cells resulted in the identification of 5,460 transcripts with at least 10 reads on average across all samples and it includes 123 peroxisomal transcripts (**Tables S1b**). For 3,952 transcripts, the respective proteins were also quantified by proteomics analysis (**Figure S1c, Table S1c**). In addition to the 91 peroxisomal transcripts/proteins detected by both methods, we found additional 32 peroxisomal transcripts which were not covered by whole cell proteomics. These include low abundance peroxisomal proteins such as the PTS2 co-receptors Pex18 and Pex21, the alternative PTS1 receptor Pex9, or the RING finger proteins Pex2 and Pex10 (**Table S1c**). Overall, 102 transcripts were regulated by a factor of at least 1.5-fold (25/77 transcripts increased/reduced; p-value < 0.05, n=3) in *pex3* cells. Remarkably, *PEX3* deletion significantly affected only five soluble peroxisomal proteins at the transcript level, which were all upregulated in *pex3* cells (**Figure 2c**).

To obtain a first overview of the main processes affected by the loss of peroxisomes in acetate-grown *S. cerevisiae* cells, we performed Gene Ontology (GO) term overrepresentation analysis for the domains “Biological Process” and “Cellular Component” for regulated proteins and transcripts with a minimum fold-change of ±1.5 and a p-value < 0.05. The analysis shows that loss of Pex3 affects proteins that are linked to “protein localization to peroxisome", “fatty acid oxidation", and, more general, "organic substance transport" and are present in the “peroxisomal membrane” including the "peroxisomal importomer complex" and the "peroxisomal matrix", whereas regulated transcripts are mainly associated with the “peroxisomal matrix” and “fatty acid beta-oxidation” (**Figure 2d**, **Table S2**).

Taken together, our proteomics data reveal that the loss of functional peroxisomes largely affects peroxisomal matrix enzymes of the β-oxidation pathway and proteins of the peroxisomal membrane. Consistent with these data, transcripts of peroxisomal β-oxidation enzymes were upregulated, whereas only non-peroxisomal transcripts were found to be downregulated in *pex3* cells. Furthermore, the regulation of numerous non-peroxisomal proteins and transcripts in *pex3* cells indicates that the elicited cellular responses go beyond peroxisomes.

### Functional classification of regulated peroxisomal proteins and transcripts in *pex3* cells

To examine the changes in *pex3* cells in detail, we classified peroxisomal proteins into different functional groups: (1) peroxisomal biogenesis factors (peroxins, *PEX* genes), (2) beta-oxidation enzymes and other peroxisomal matrix proteins, (3) proteins with a function in the glyoxylate cycle, and (4) solute transporters. In our analysis of proteomics and transcriptomics data displayed in volcano plots, we also directly compared changes in the proteome and transcriptome in consequence of *PEX3* deletion. To this end, the mean log_2_ *pex3*/WT ratios of transcripts and proteins were plotted against each other.

### Changes in the levels of peroxins are independent of transcription

Peroxisomal proteins involved in the biogenesis of peroxisomes (peroxins, *PEX* genes) are of crucial importance for peroxisomal functions, and in *S. cerevisiae,* 29 peroxins have been identified so far (Jansen et al., 2021). In total, we quantified 19 peroxins at the protein level and all *PEX* transcripts except for *PEX3*. We found that membrane-localized peroxins, which play a role in peroxisomal matrix protein import, are considerably decreased in abundance at the whole cell level (**Figure 3a**). Exceptions are Pex8, which is attached to the luminal side of the peroxisomal membrane (Rehling et al., 2000), and Pex22. The members of the Pex30 family, Pex29, Pex30, and Pex31, which do not reside in the peroxisomal membrane, but in the cortical ER membrane (David et al., 2013; Mast et al., 2016), were also unchanged. The opposite regulation was found for the cytosolic receptor proteins Pex5 and Pex7, which are essential for the import of PTS1- and PTS2-containing proteins, respectively. The finding was most pronounced for the PTS1 receptor Pex5, with a 2.7-fold increase in abundance.

**Figure 3.**
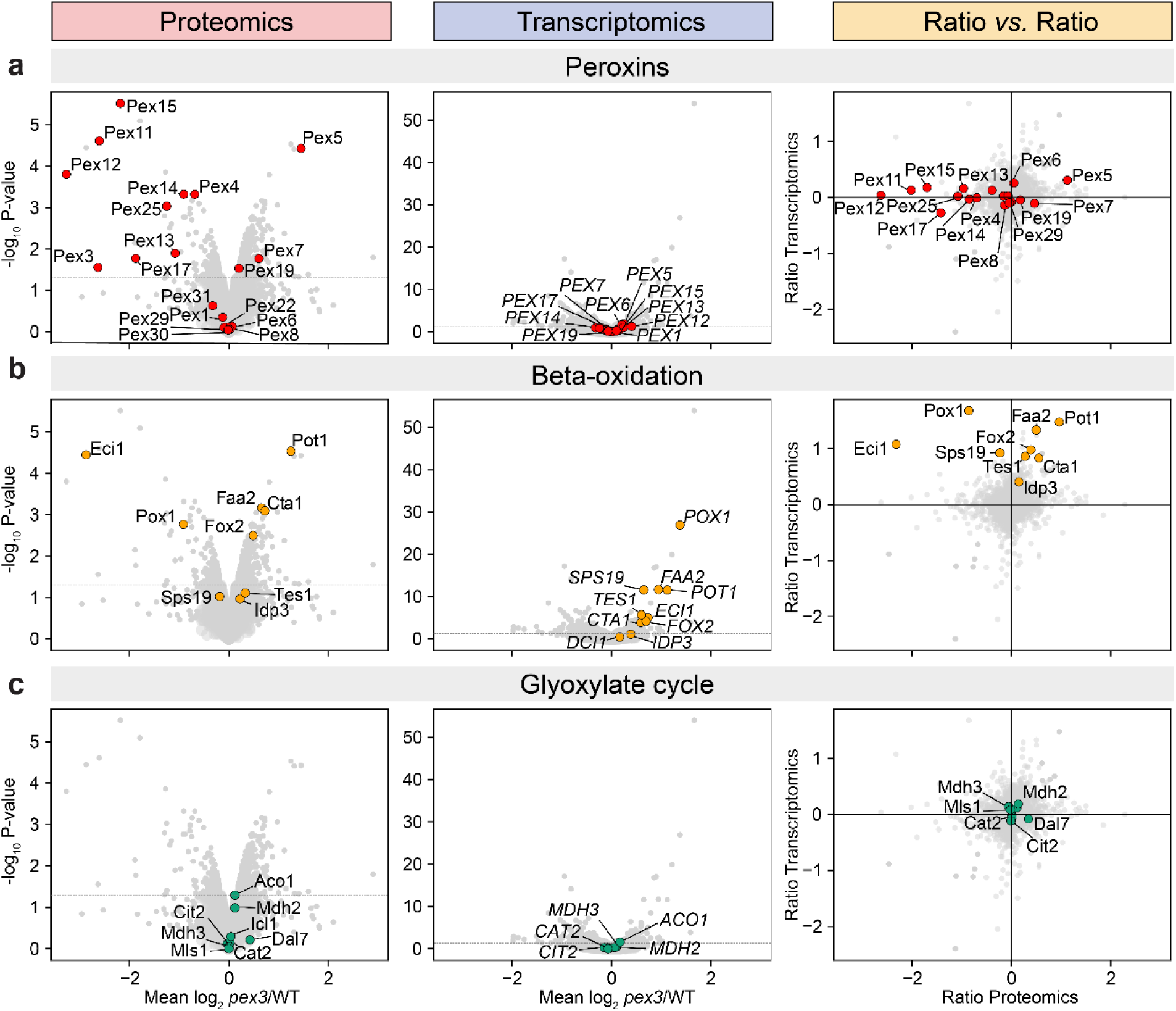
Effect of the loss of Pex3 on the levels of proteins and transcripts associated with peroxisomal processes. Same volcano plots as in Figures 2b and 2c depicting the results of the quantitative proteomics (left) and transcriptomics (middle) analyses and ratio-*versus*-ratio plots of both omics data (right) highlighting all peroxins (a) as well as proteins/transcripts related to beta-oxidation (b) and the glyoxylate cycle (c). The horizontal line in the volcano plots indicates a p-value of 0.05; the horizontal and vertical lines in the ratio-*versus*-ratio plots indicate a log2-fold change of 0.

The AAA-proteins Pex1 and Pex6 are essential for the peroxisomal export of the Pex5 to the cytosol (Platta et al., 2005). Interestingly, both peroxins were not altered in abundance, although levels of their peroxisomal membrane anchor Pex15 were strongly decreased. This is likely related to the fact that Pex1 and Pex6 can form a stable, soluble hexameric complex in the absence of Pex15 (Saffian et al., 2012). We observed an opposite regulation for Pex4, which is involved in Pex5 ubiquitination after the translocation of cargo proteins into the peroxisomal matrix (Ali et al., 2018; Williams et al., 2012). Whereas levels of its membrane anchor Pex22 remained unchanged, Pex4 abundance was considerably decreased in the absence of Pex3 (**Figure 3a**).

In *S. cerevisiae,* Pex3 and Pex19 have been identified as necessary for PMP targeting (Hettema et al., 2000). In the absence of Pex3, we found that the cytosolic PMP receptor/chaperone Pex19 was only slightly increased in abundance, whereas levels of Pex19 cargo PMPs were reduced, including peroxins involved in (i) the import of peroxisomal matrix proteins [Pex14, Pex13, Pex17 (docking complex), Pex12 (RING complex), and Pex15 (export complex)], and (ii) peroxisome proliferation (Pex11 and Pex25). In stark contrast to these proteome data, transcriptomics analysis revealed that none of the *PEX* transcripts were regulated in *pex3* cells (**Figure 3a, Table S1d)**.

Other proteins that are relevant in peroxisome biology, such as proteins that play a role in organelle fission (Dnm1, Vps1), inheritance (Inp1, Inp2) or pexophagy (Atg36) were all unchanged at the protein and transcript level (**Table S1d**).

To conclude, our omics data indicate that *PEX3* deletion results in unbalanced levels of peroxins involved in the late steps of peroxisomal matrix protein import, which involves the recycling of the peroxisomal receptor proteins Pex5 and Pex7. Both cytosolic receptor proteins accumulate in the absence of Pex3, which might result from the reduction in ubiquitination factors, Pex4 and Pex12. Since transcript levels remained unaltered in *pex3* cells, the observed changes in peroxin abundances are governed by posttranscriptional mechanisms, with membrane-localized peroxins presumably degraded in the absence of the target membrane.

### Central β-oxidation enzymes are upregulated in the absence of Pex3

We asked whether the increased abundance of the PTS1 and PTS2 receptor proteins is accompanied by increased levels of peroxisomal matrix proteins. To address this, we examined changes in protein and transcript levels of peroxisomal enzymes in *pex3* compared to WT cells. Remarkably, β-oxidation enzymes (i.e., Fox2, Pot1, Faa2) were consistently upregulated at protein and transcript level (**Figure 3b**; **Table S1d**), with a few exceptions: first, auxiliary β-oxidation enzymes (Tes1, Sps19) and peroxisomal isocitrate dehydrogenase (Idp3) that is required for β-oxidation were mildly affected only at transcript level, and second, Pox1 and Eci1 were markedly decreased at protein but increased at transcript level. Recent studies revealed that the protein levels of Pox1 are posttranslationally regulated by Arf1 (Enkleret al., 2023). Possibly, this process is responsible for the low Pox1 levels in *pex3* cells.

Pox1 produces hydrogen peroxide during β-oxidation, which is degraded by peroxisomal catalase (Cta1) (Hiltunen et al., 2003). In *pex3* cells, we found Cta1 to be upregulated at transcript and protein levels (**Figure 3b**). The peroxisomal Δ3-cis-Δ2-trans-enoyl-CoA isomerase Eci1 is required for the degradation of unsaturated fatty acids (Gurvitz et al., 1998). Previous work showed that Eci1 is co-imported with the Δ^3,5^-Δ^2,4^-dienoyl-CoA isomerase Dci1 (Yang et al., 2001), an enzyme involved in peroxisomal β-oxidation (Gurvitz et al., 1999). Since transcripts of its partner protein Dci1 (not identified on protein level) were not increased, Eci1 protein alone might not be stable and is therefore degraded in *pex3* cells (**Figure 3b**). In addition to the above mentioned β-oxidation enzymes, which all contain a PTS1, the essential PTS2 protein 3-ketoacyl-CoA thiolase (Pot1), which catalyzes the last step of each β-oxidation cycle, was upregulated to a similar extent at protein and transcript level (**Figure 3b**). Further upregulated peroxisomal enzymes were Lpx1 (a lipase) and Pcs60 (involved in oxalate degradation), but others (i.e. Bud16, Lys1, Str3) were unchanged (**Figure S2**). Interestingly, protein levels of the stress inducible peroxisomal enzymes Gpd1 (glycerol phosphate dehydrogenase) and Pnc1 (nicotinamidas) also not increased (Choudhry et al., 2016; Effelsberg et al., 2015), but both were transcriptionally downregulated (**Figure S2**).

To underscore our findings on the specific upregulation of β-oxidation enzymes, we also examined proteins of the glyoxylate cycle, which is another common and conserved function of yeast peroxisomes besides β-oxidation. Independent of their peroxisomal or cytosolic localization, transcripts and protein levels of the known glyoxylate cycle enzymes were not affected by the deletion of *PEX3* (**Figure 3c**).

To conclude, in the absence of Pex3, central peroxisomal β-oxidation enzymes are upregulated at transcript and protein levels in acetate grown yeast cells. Exceptions are Eci1 and Pox1 exhibiting increased transcript but reduced protein levels in *pex3* cells, which involves posttranslational mechanisms for their selective removal in peroxisome-depleted cells. The upregulation of β-oxidation enzymes might be linked to the increased abundance of the receptor proteins Pex5 and Pex7 and/or may be a direct cellular response to counteract the loss of functional peroxisome.

### Mpc1 and Mpc3 are so far unknown transporters of the peroxisomal membrane

Peroxisomal functions including the degradation of fatty acids require the transport of ATP, nucleotides, and metabolites across the peroxisomal membrane. Thus, based on the metabolic pathways that occur in peroxisomes, numerous solute transporters are predicted to exist in the peroxisomal membrane, but only a few have been identified so far. We found that levels of the peroxisomal adenine nucleotide transporter Ant1, which is involved in β-oxidation (Palmieri et al., 2001; van Roermund et al., 2001) were markedly decreased at the protein but not transcript level in *pex3* cells (**Figure 4a**). Other known peroxisomal transporters involved in the import of activated long-chain fatty acids (Pxa1/2) and oligopeptides (Opt2) were only detected at the transcript level, without significant changes in *pex3* cells (**Figure 4a**; **Table S1d**).

**Figure 4.**
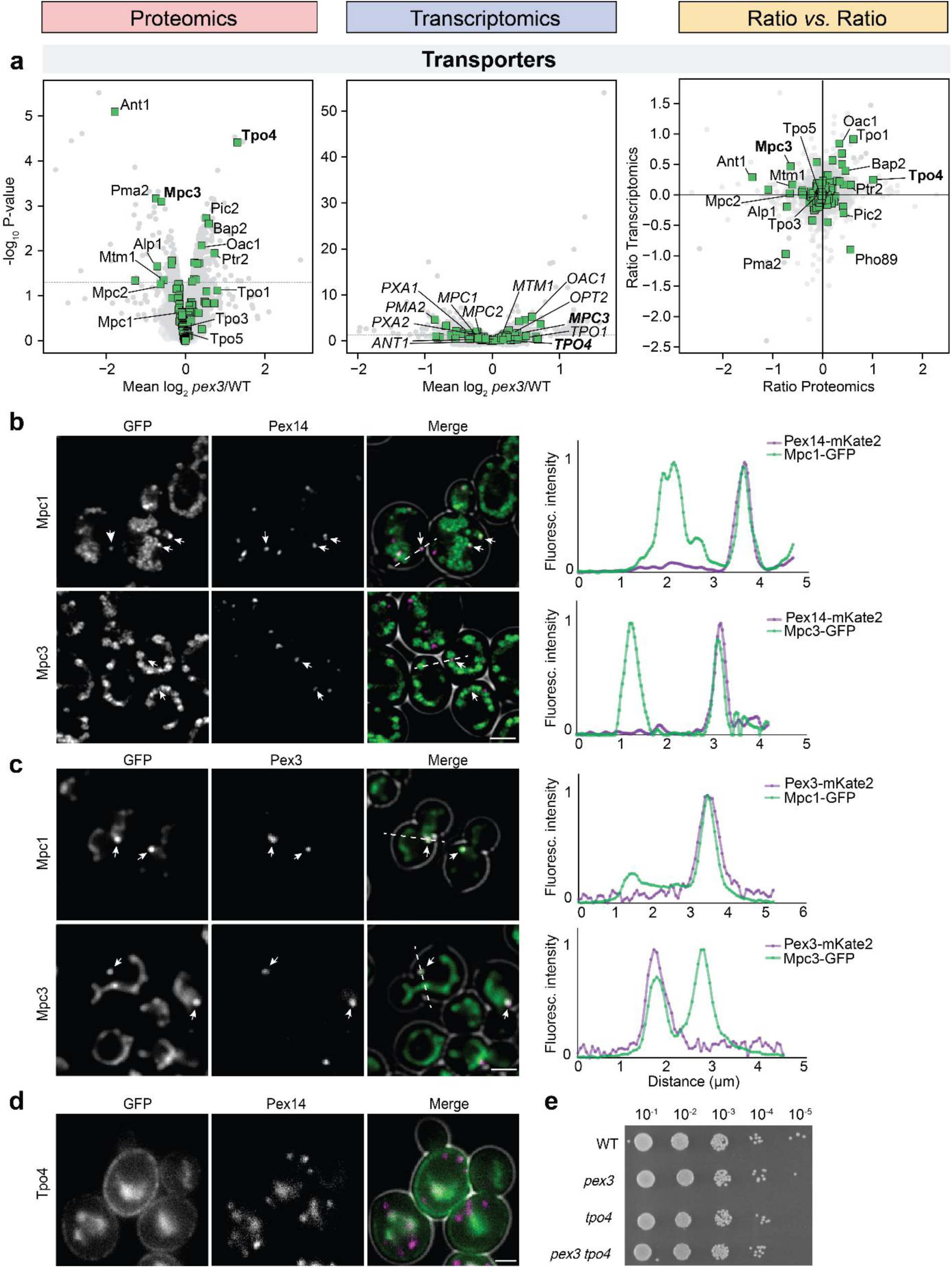
Dual localization of Mpc proteins in yeast. (a) Abundance of transporter proteins and transcripts in *pex3 versus* WT cells. The same plots as in Figure 3 highlighting transporters. (b) FM analysis of acetate grown *S. cerevisiae* cells producing Mpc1-GFP or Mpc3-GFP under the control of their endogenous promoters. Pex14-mCherry was used as a peroxisome marker. Scale bar: 2.5 μm. Graphs show normalized fluorescence intensity along the lines indicated in the merged images. (c) FM analysis of glucose grown *H. polymorpha* cells producing Mpc1-GFP and Mpc3-GFP under the control of the *ADH1* promoter. Pex3-mKate2 was used as a peroxisome marker. Scale bar: 2.5 μm. Graphs show normalized fluorescence intensity along the lines indicated in the merged images. (d) FM analysis of acetate grown *S. cerevisiae* cells producing Tpo4-GFP under the control of its endogenous promoter. Pex14 mCherry was used as a peroxisome marker. Scale bar: 1.5 μm. (d) Growth analysis of WT and the indicated (double) deletion strains. Spot assays were performed using a mineral medium containing 0.5% acetate. Cultures were serially diluted (from 10^-1^ to 10^-5^) and spotted on agar plates. A representative spot assay of 2 independent experiments is shown.

Recent studies on *S. cerevisiae* peroxisomes (Yifrach et al., 2022) and mitochondria (Morgenstern et al., 2017) showed that the proteomes of both organelles contain numerous multi-localized proteins. Thus, we hypothesized that solute transporters of other cellular compartments might partially localize to peroxisomes. The assumption that only a small fraction of these transporters reside in peroxisomal membranes would make it plausible that they have not been localized in peroxisomes so far. To address this issue, we examined changes in protein and transcript levels of non-peroxisomal transporters in *pex3* cells (**Figure 4a**). Since known PMPs including Ant1 showed a significant decrease in protein but not transcript levels in *pex3* cells (**Figures 3a** and **4a**), we searched for proteins with similar behavior. This revealed two mitochondrial inner membrane carriers, Mtm1 and Mpc3. Mtm1 is a likely candidate for a peroxisomal transporter because it transports pyridoxal 5’-phosphate, which is produced by the putative pyridoxal kinase Bud16, an enzyme that localizes to the cytosol and peroxisomes (Yifrach et al., 2022). Unfortunately, we were unable to detect Mtm1-GFP produced under the control of its endogenous promoter by FM. We next tested the pyruvate carrier Mpc3 (Bricker et al., 2012) (**Figure 4a**). Importantly, pyruvate is produced in yeast peroxisomes by the matrix enzyme Str3. Therefore, a pyruvate transporter is predicted to occur in the peroxisomal membrane. In respiring cells, Mpc3 forms a heterodimer with Mpc1 (Tavoulari et al., 2019), whereas Mpc2 heterodimerizes with Mpc1 to form the fermentative mitochondrial pyruvate carrier (MPC) isoform (Bender et al., 2015; Herzig et al., 2012). In our omics data, we quantified Mpc1, Mpc2, and Mpc3, with Mpc3 being significantly decreased in protein abundance (**Figure 4a**; **Table S1d**). Notably, Str3 levels were also reduced in *pex3* cells (**Figure S2**). Since Mpc1 is part of both MPC isoforms, a minor decrease in abundance due to the loss of peroxisomes might not be detectable in our experimental setting. To further test whether a portion of Mpc1 and Mpc3 localizes to peroxisomes in addition to their main localization in mitochondria, we performed FM analysis using proteins C-terminally tagged with GFP. As shown in **Figure 4b**, Mpc1 and Mpc3 predominantly localize to mitochondria, but a small fraction of these proteins is also present in small spots that harbor the peroxisomal marker protein Pex14.

Complementary to *S. cerevisiae, H. polymorpha* is a methylotrophic yeast that is extensively used as a model organism in peroxisome research. *H. polymorpha* has two Mpc proteins, here called *Hp*Mpc1 (80.2 % similarity with *Sc*Mpc1) and *Hp*Mpc3 (73.4 % similarity to *Sc*Mpc3). To analyze whether the localization of a small portion of the Mpc proteins on peroxisomes also occurs in *H. polymorpha*, we also checked the localization of the two *H. polymorpha* Mpc proteins. FM analysis showed that like in *S. cerevisiae,* both proteins have a mitochondrial and peroxisomal localization (**Figure 4c**). These results imply that the dual localization of Mpc proteins is conserved in different yeast species.

In contrast to Ant1, Mtm1 and Mpc3, we also detected transporters of increased protein abundance in *pex3* cells (**Figure 4a**). Tpo4 was one of the proteins with the highest change in abundance in *pex3* cells. Tpo1-4 are members of the major facilitator superfamily, localized in the vacuole- and plasma membrane. We also detected Tpo1, 2 and 3, but only Tpo4 was significantly enhanced at the protein level. Tpo transporters are implicated in transport of various compounds, including drugs (Tpo1) (Albertsen et al., 2003; do Valle Matta et al., 2001), polyamines (Tpo1-4) (Tomitori et al., 2001, 1999), acetate (Tpo2 and Tpo3) (Zhang et al., 2022) and medium chain fatty acids (Tpo1) (Hu et al., 2020). Tpo1,2 and 3 have been extensively studied, however, Tpo4 is poorly characterized. FM analysis of a strain producing Tpo4-GFP revealed that the protein was predominantly present at the plasma membrane, but a portion of the protein localizes to the vacuole (**Figure 4d**). Tpo4-GFP did not co-localize with the peroxisomal marker protein Pex14. The enhanced levels of Tpo4 in *pex3* cells imply that it may be involved in the secretion of potentially toxic compounds that might accumulate in the cell in the absence of peroxisomes. To investigate the correlation between Tpo4 and peroxisomes, we analyzed growth on acetate using a spot assay. As shown in **Figure 4e**, the *tpo4* single and *tpo4 pex3* double deletion strains showed similar growth. This indicates that enhanced Tpo4 levels are not crucial for the growth of *pex3* cells on acetate.

In sum, we investigated transporters with significant changes at the protein level. While for Tpo4 we did not find any direct correlation with peroxisomes, our investigation of the pyruvate carriers Mpc1 and Mpc3 revealed that a portion of these proteins is detectable at some of the peroxisomes.

### Functional classification of regulated non-peroxisomal transcripts and proteins in *pex3* cells

Our omics analysis of *pex3* cells revealed that peroxisomal membrane-localized peroxins (except for Pex22 and Pex8) were decreased in abundance. Previous work showed that Pex11 is detectable on mitochondria in yeast *pex3* and/or *pex19* cells (Nuebel et al., 2021; Wróblewska et al., 2017). Thus, in the absence of functional peroxisomes, PMPs are degraded and/or relocalized to another subcellular niche. In yeast, all peroxisomal proteins are synthesized in their mature form and folded in the cytosol prior to import. However, the upregulation of abundant peroxisomal β-oxidation enzymes might challenge cellular proteostasis and the loss of peroxisomal functions might elicit metabolic changes. To obtain information about the existence of such cellular responses, we further investigated changes in the abundance of non-peroxisomal transcripts and proteins in *pex3* cells.

### Pex3 deficiency results in altered levels of various factors of non-peroxisomal metabolic pathways

We next assessed changes in the levels of non-peroxisomal transcripts and proteins in *pex3* cells (**Figure 5**). Many of the regulated non-peroxisomal transcripts that encode proteins whose levels were unchanged are uncharacterized ones (**Table S1d**), and that is why we do not discuss them further. Among the characterized regulated transcripts, we found a strong transcriptional upregulation of *EEB1*, which is involved in the biosynthesis of medium-chain fatty acid ethyl ester under fermentative growth (Saerens et al., 2006), and two helicases (YBL113C, YBL111C), but none of their translation products was identified in our proteomics analysis (**Figure 5a**; **Table S1d**). Upregulated transcripts that displayed a slight increase at the protein level include the cytosolic proteins inositol-3-phosphate synthase (Ino1) and Leu1, which is essential for leucine biosynthesis, as well as the branched-chain amino acid aminotransferase Bat1, and the oxaloacetate carrier Oac1 (**Figure 5a**).

**Figure 5.**
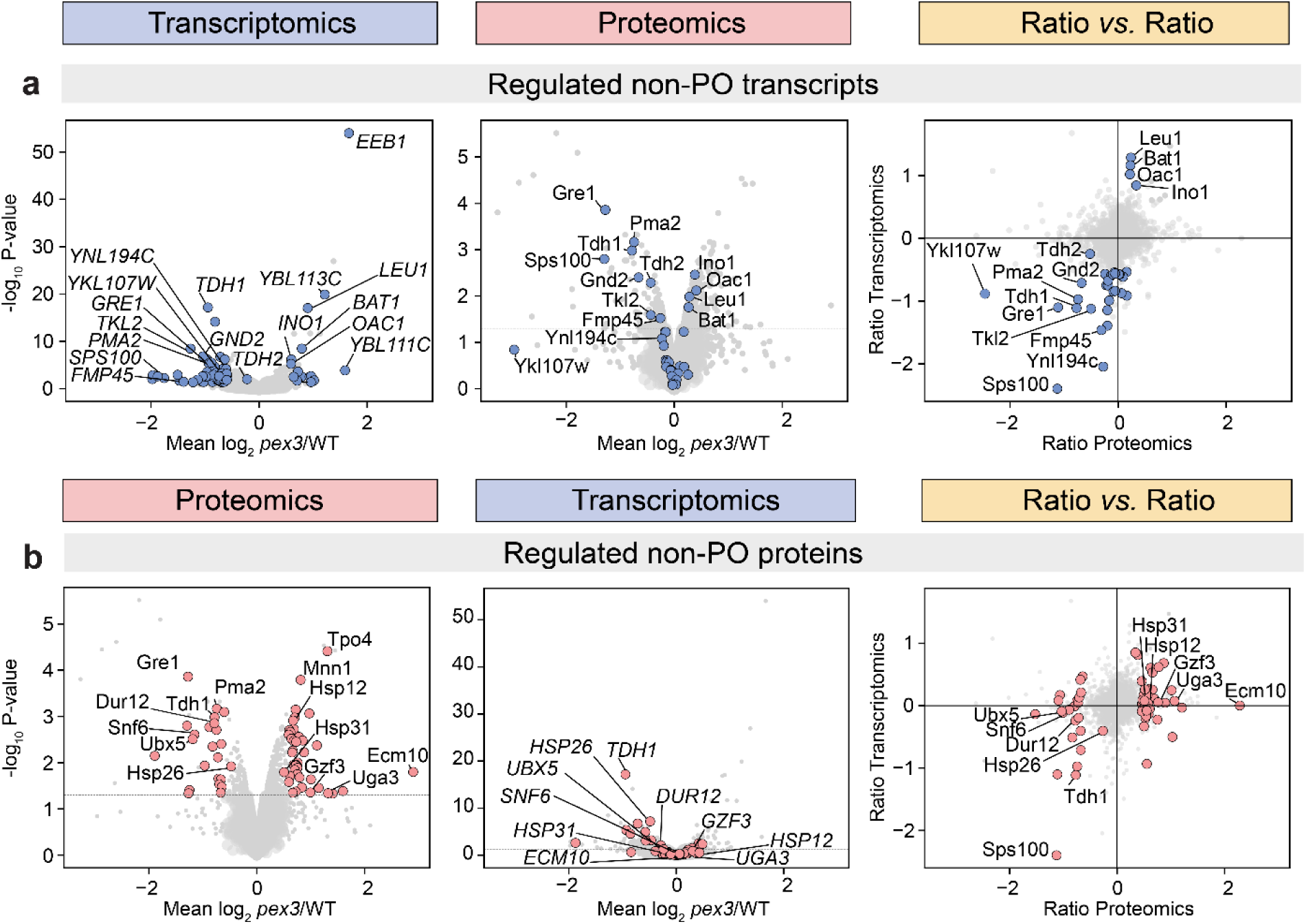
Effect of *PEX3* deletion on transcripts and proteins without a link to peroxisomal processes. Same plots as in Figure 3. Highlighted are (a) regulated transcripts unrelated to peroxisomal (PO) processes and the corresponding proteins and (b), *vice versa*, regulated non-peroxisomal proteins and the corresponding transcripts. Transcripts/proteins with a minimum fold-change of ± 1.5 and a p-value of < 0.05 (n=3) are highlighted in blue/red, as well as the TDH2 transcript.

Among the downregulated transcripts that were also reduced at protein level were the plasma membrane H^+^-ATPase Pma2, an aldehyde reductase (Ykl107w), glyceraldehyde-3-phosphate dehydrogenases (Tdh1; Tdh2, 1.3-fold up), a spore wall maturation associated protein (Sps100), the general stress responsive factor Gre1, as well as the pentose phosphate pathway enzymes 6-phosphogluconate dehydrogenase (Gnd2) and transketolase (Tkl2). Furthermore, transcripts of the plasma membrane protein Ynl194c and its paralog Fmp45, which both are involved in sphingolipid metabolism (Young et al., 2002), were considerably downregulated (**Figure 5a**).

We next examined regulated non-peroxisomal proteins, which were in many cases not affected at the transcript level in *pex3* cells (**Figure 5b**). Interestingly, this included a few proteins related to nitrogen metabolism (Uga3, Gzf3, Dur12). Both Uga3 and Gzf3 are enhanced at the protein level. Uga3 and Gzf3 are transcription factors involved in the regulation of genes involved in the uptake and utilization of non-preferred nitrogen sources (Palavecino-Ruiz et al., 2017). One of the Gzf3 regulated genes is urea amidolyase Dur12 (alias Dur1,2) (Chisholm and Cooper, 1984; Coulon et al., 2006; Wu et al., 2020), which had a lower abundance in the *pex3* cells. Since ammonium is the nitrogen source in the medium used, it is not clear why proteins involved in the utilization of alternative nitrogen sources were changed.

### Loss of Pex3 leads to increased levels of specific chaperones ensuring protein folding

Among the proteins of decreased abundance were Ubx5, Snf6, and Hsp26. Ubx5 is an adapter of the AAA ATPase Cdc48/p97 (Schuberth et al., 2004) and involved in the degradation of proteins by the proteasome (Li et al., 2023 *preprint;* Noireterre et al., 2023; Verma et al., 2011). The downregulation of the nuclear protein Snf6, which is involved in transcriptional activation (Estruch and Carlson, 1990; Laurent et al., 1991), generally fits the observation that *pex3* cells do not show widespread changes in their transcriptional program. The small heat shock protein Hsp26 is involved in the reactivation of misfolded, aggregated proteins by Hsp104 (Cashikar et al., 2006). Like the stress factor Gre1, Hsp26 was transcriptionally downregulated (**Figure 5b**). Furthermore, *PEX3* deletion resulted in decreased transcripts but unaltered protein levels of the disaggregase Hsp104 and the sequestrase Hsp42 (**Table S1d**), which both fulfill vital roles in maintaining proteostasis in *S. cerevisiae* (Bösl et al., 2006; Krämer et al., 2023; Specht et al., 2011). However, we found an increase in the abundance of Hsp12 and Hsp31 (1.4-fold up) (**Figure 5b**). Hsp12 is involved in plasma membrane organization during stress (Sales et al., 2000; Welker et al., 2010), whereas Hsp31 protects cells from protein misfolding stress at an early state to prevent protein aggregation (Tsai et al., 2015). Thus, our data indicate that early protective responses to mitigate protein misfolding are active and enable cells to maintain proteostasis in the absence of peroxisomes.

Most interestingly, we observed the highest increase in abundance for the mitochondrial matrix protein Ecm10 in *pex3* cells, but transcripts remained unchanged (**Figure 5b**). Ecm10 is a member of the mitochondrial Hsp70 protein family. Further family members are Ssc1 and Ssq1 (reviewed in Voos and Röttgers, 2002). Ssc1 is involved in the import and folding of mitochondrial precursor proteins and Ssq1 in iron-sulfur protein biogenesis, but neither of them was regulated in *pex3* cells. Other mitochondrial chaperones (e.g. Hsp60, Hsp78, Hsp10) were also not changed in abundance (**Table S1d**). Notably, Ecm10 is a paralog of Ssc1, and supposedly also involved in the import and folding of newly synthesized precursor proteins in the mitochondrial matrix (Baumann et al., 2000). Based on this unexpected specific mitochondrial response manifested by highly increased Ecm10 levels, we examined in more detail changes in the abundance of mitochondrial proteins in *pex3* cells which lack functional peroxisomes.

### Proteins with a central role in mitochondrial respiration and import are increased in *pex3* cells

Mitochondria fulfill central metabolic functions highlighted by the production of ATP by oxidative phosphorylation (Morgenstern et al., 2017; Pfanner et al., 2019). In *pex3* cells, most mitochondrial transcripts were unaltered in abundance, except for the downregulated transcripts of the Sdh3 and Sdh4 homologs Shh3 and Shh4 or upregulated transcripts of the oxaloacetate carrier protein Oac1 and mitochondrial aldehyde dehydrogenase Ald5, which plays a role in the biogenesis of the electron transport chain and synthesis of acetate (Kurita, 1999; Saint-Prix et al., 2004) (**Figure S3a**). On the proteome level, we found an increase in the abundance of numerous proteins that are associated with central mitochondrial protein machineries in *pex3* cells. We found a higher abundance of proteins with a role in the assembly and function of mitochondrial respiratory chain complexes (**Figure 6**). These included (i) different subunits (Atp18, Atp15, Atp14) of the mitochondrial F1F0 ATP synthase, (ii) the twin ATPase regulators Inh1 and Stf1 that inhibit uncoupled ATP hydrolysis (Venard et al., 2003), (iii) assembly factors (Cox19, Cox17, Cox12) of mitochondrial cytochrome c oxidase (Das et al., 2021; Glerum et al., 1996; Nobrega et al., 2002; Rigby et al., 2007), and (iv) cytochrome c isoform 1 (Cyc1) and cytochrome c heme lyase Cyc3, which attaches heme to apo-Cyc1 (Dumont et al., 1991).

**Figure 6.**
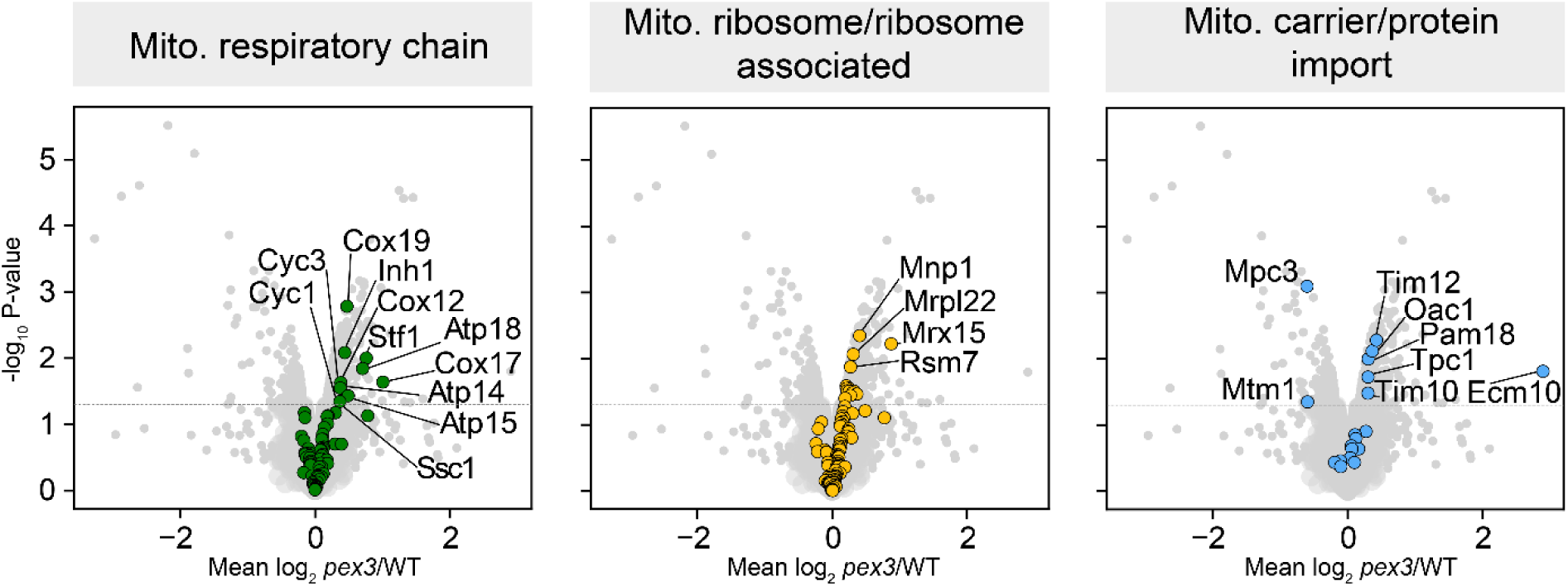
Changes in the abundance of mitochondrial proteins in *pex3* cells. The same volcano plots as in Figure 2b high lighting components of the mitochondrial (mito.) respiratory chain (left), mitochondrial ribosomal and ribosome-associated components (middle), and proteins associated with the mitochondrial carrier import pathway or involved in the mitochondrial presequence import pathway (right).

The mitochondrial respiratory chain is of dual-genetic origin and therefore its assembly requires the insertion of nucleus- and mitochondria-encoded subunits into the inner mitochondrial membrane (IMM). Interestingly, *pex3* cells exhibited higher levels of Dpi29 (Morgenstern et al., 2017), alias Mrx15 (**Figure 6**). It is a receptor of mitochondrial ribosomes required for the efficient co-translational insertion of mitochondria-encoded membrane proteins into the IMM (Möller-Hergt et al., 2018). Moreover, proteins of mitochondrial ribosomes were generally shifted towards higher abundance in *pex3* cells, which was most pronounced for Mnp1, Mrpl22 or Rsm7 (**Figure 6**). In addition to mitochondria-encoded proteins, also nucleus-encoded hydrophobic metabolite carriers need specific guidance for their delivery to the carrier translocase TIM22 in the IMM (Pfanner et al., 2019). In *pex3* cells, the abundance of the mitochondrial carrier proteins Oac1 and Tpc1 was increased, along with higher levels of Tim10 and Tim12, which deliver, together with Tim9, carrier proteins to the carrier translocase TIM22 in the IMM (**Figure 6**). Levels of the J-protein cochaperone Pam18 were also increased, which is an essential subunit of the presequence-associated motor of the translocase of the inner mitochondrial membrane (TIM23) (Pfanner et al., 2019). Specifically, Pam18 facilitates the import of mitochondrial presequence proteins into the mitochondrial matrix by stimulating the ATPase activity of the Hsp70 family ATPase Ssc1 (Truscott et al., 2003). Notably, levels of Ecm10, but not Ssc1, were highly increased, which points to a role of Ecm10 in mitochondrial protein import under peroxisome-deficient conditions. Moreover, the abundance of the integral ER membrane protein Ylr050C, alias Ema19, and the intramembrane aspartyl protease of the perinuclear ER membrane (Ypf1) were also increased in cells lacking Pex3 (**Figure 3b**). Ema19 is involved in the ER surface retrieval pathway (ER-SURF) to effectively target membrane proteins to the mitochondrial surface for import (Hansen et al., 2018) and has been identified as a quality control factor for newly synthesized but unstable mitochondrial precursor proteins (Laborenz et al., 2021). Ypf1 is a protease that is involved in a branch of the endoplasmic reticulum unfolded protein response (ERAD) (Avci et al., 2014).

To conclude, our omics data show that the absence of Pex3 acetate grown yeast cells does not elicit major changes in the transcriptional program and changes in protein levels are therefore mainly governed by posttranscriptional mechanisms. Our proteomics data further indicate that the loss of functional peroxisomes leads to changes in the abundance of specific factors that play important roles in mitochondrial respiration, protein import and the folding of mitochondrial precursors in the mitochondrial matrix.

## Discussion

Here, we report a comprehensive whole cell transcriptome and proteome study to uncover the effects of the absence of peroxisomes in yeast cells by comparing WT and peroxisome-deficient *pex3* cells. We show that the absence of functional peroxisomes elicits specific cellular responses (for an overview see **Figure 7**). Key observations include the upregulation of almost all genes encoding enzymes of the peroxisomal β-oxidation pathway, together with enhanced levels of the corresponding proteins. In contrast, the expression of *PEX* genes was unchanged at the transcript level, while most peroxins were reduced (all peroxisomal membrane bound peroxins) at the protein level. Exceptions were the receptor proteins Pex5 and Pex7, which were considerably increased in abundance. Intriguingly, none of the peroxisome related genes, but several genes encoding proteins involved in various other cellular processes, were downregulated in *pex3* cells. These include proteins of the pentose phosphate pathway (PPP) and plasma membrane transporters. Moreover, specific mitochondrial proteins were enhanced, although the corresponding transcripts were unchanged (**Figure 7**).

**Figure 7.**
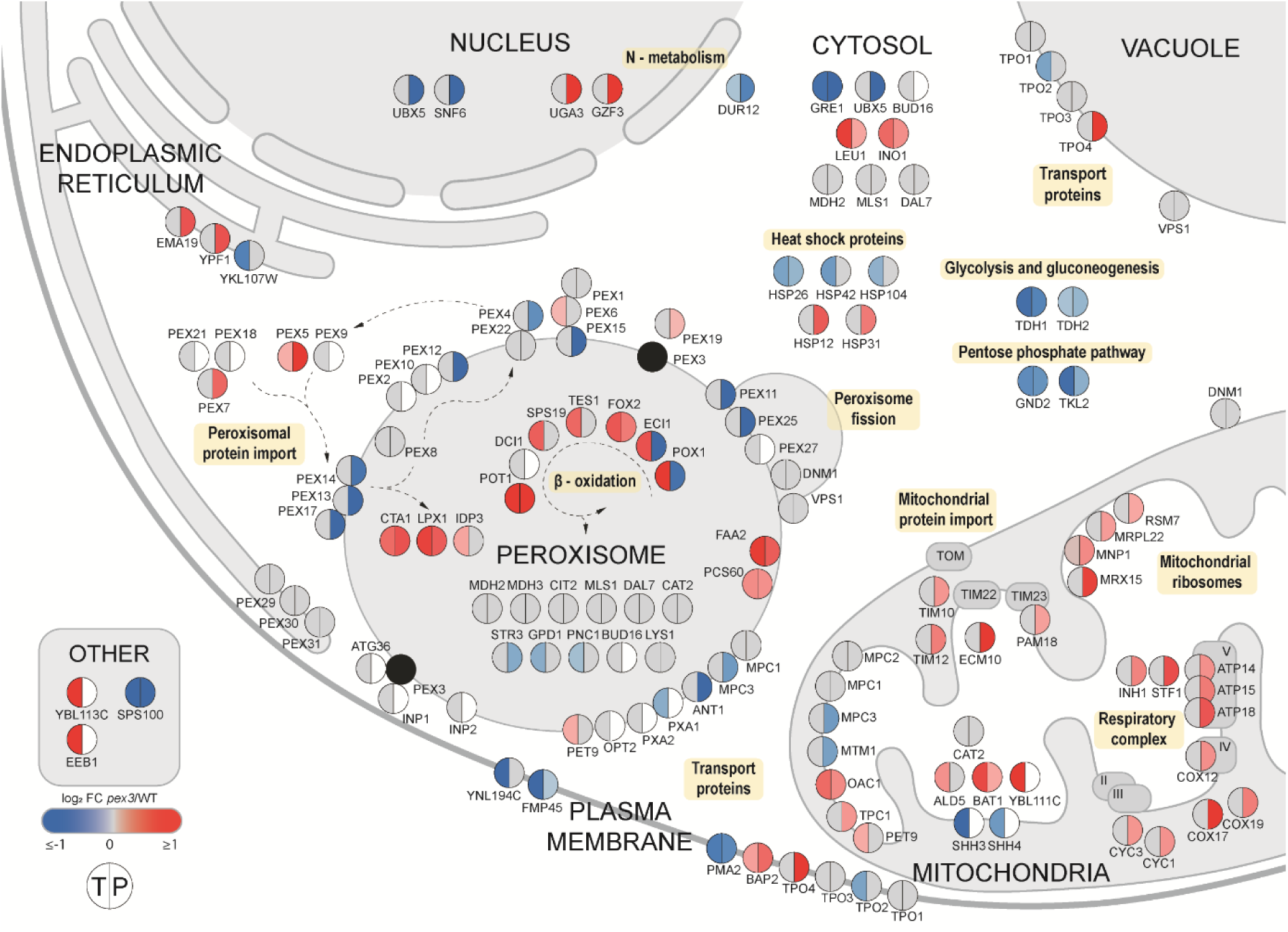
Overview of changes in the transcriptome and proteome of *pex3* versus WT cells. Shown are transcripts/proteins discussed in the results section and their subcellular localization (based on literature, the *Saccharomyces* Genome Database and/or UniProt). Circles represent the transcripts (T, left) and proteins (P, right). Transcripts/proteins that fulfill the p-value threshold of < 0.05 are colored. Unchanged and undetected transcripts/proteins are depicted in gray and white, respectively. Category "other" includes candidates with unknown localization.

We used yeast *pex3* cells, which lack functional peroxisomes, as a model system. Peroxisome-deficient yeast mutants are viable (Erdmann et al., 1989), while the complete absence of any other cell organelle is lethal in yeast. This unique feature was the basis of our study. To detect changes that were exclusively caused by the absence of peroxisomes, it was imperative to use conditions that do not affect cell growth or repress genes encoding proteins involved in peroxisome biology. We showed that growth on acetate fulfils these criteria. In addition, under these conditions, peroxisomes harbor enzymes of the glyoxylate cycle (Kunze et al., 2006), which are essential for the growth on acetate (Lee et al., 2011; Weill et al., 2018).

Our data show that the levels of the transcripts and proteins of the glyoxylate cycle enzymes were unchanged in *pex3* cells. Apparently, these cytosolically mislocalized enzymes are sufficiently active to allow growth on acetate. This is in agreement with the early observation that *S. cerevisiae pex* mutants are capable of growing on non-fermentable carbon sources such as acetate and only are defective in the utilization of oleic acid (Erdmann et al., 1989).

Despite the absence of peroxisomes in *pex3* cells, the transcriptome hardly changed. Only 102 out of 5,460 detected transcripts were altered more than ±1.5-fold (p-value < 0.05), which is less than 2%. Also, only 17 out of the 123 transcripts encoding peroxisomal proteins had a p-value of < 0.05, with 9 showing a fold change of > ±1.5-fold. Similar observations were made in transcriptomic studies using other *S. cerevisiae pex* mutants (*pex12*; microarray analysis (Breitling et al., 2002) or *pex19*; RNAseq (Nuebel et al., 2021). Notably, for *S. cerevisiae pex12* a more than 1.5-fold upregulation of seven genes involved in lysine biosynthesis was reported. We did not observe changes in any of these transcripts in *S. cerevisiae pex3*. Most likely, this is due to differences in growth conditions (glucose instead of acetate, i.e. fermentative versus non-fermentative conditions) or the method used (RNA-seq versus microarrays). Nuebel and colleagues observed that three zinc-response genes (*ADH4, ZAP1, ZRT1*) were downregulated in *S. cerevisiae pex19* cells. The transcripts of none of these genes were significantly changed in our study using acetate grown *pex3* cells. In *pex19* cells, *ULI1* transcripts were reduced as well (Nuebel et al., 2021). Uli1 is a protein of unknown function that is induced by ERAD. The ERAD pathway targets misfolded ER proteins for retrotranslocation into the cytosol and degradation via the ubiquitin-proteasome system (Hiller et al., 1996; Krshnan et al., 2022; Ward et al., 1995). In *pex3* cells, we found higher levels of the protease Ypf1, which is involved in the selective degradation of functional proteins as part of a distinct branch of ERAD and was shown to regulate the levels of nutrient transporters during starvation (Avci et al., 2014). Thus, it might be involved in regulating protein levels in yeast cells lacking Pex3.

Another ERAD related protein that was altered in our study is Ubx5, an adaptor of the Cdc48 machine in ERAD (Schuberth et al., 2004). In the study using *pex19* cells by Nuebel and colleagues, the growth conditions were quite different from what we used (i.e. growth on a mixture of ethanol and glycerol and harvesting of cells from the stationary growth phase), which may explain the differences in observations.

Like for transcripts, also a rather small fraction of the entire proteome was changed in *pex3* cells. Out of the 92 detected peroxisomal proteins, 17/11 were reduced/increased in abundance (p-value < 0.05). Strikingly, almost all known PMPs, which include peroxins and the peroxisomal transporter Ant1, had a reduced abundance in *pex3* cells. This is in agreement with earlier studies, which revealed that newly synthesized PMPs were rapidly degraded in *S. cerevisiae pex3* cells (Hettema et al., 2000). The instability is most likely due to the fact that PMP sorting is abolished. Previous FM studies revealed that *PEX3* deletion results in the mislocalization of Pex11 to mitochondria in oleic acid induced yeast cells (Motley et al., 2015; Wróblewska et al., 2017). Under these conditions, Pex14 and Pex8 were detected at peroxisomal membrane vesicles, while the levels of the other PMPs studied (Pex10, Pex13, Ant1) were too low to determine their localization by FM (Wróblewska et al., 2017). Hence, in the absence of Pex3, the bulk of the newly synthesized PMPs is degraded, while a portion of some of them localizes to membrane vesicles or mitochondria.

Interestingly, like for PMPs, the protein level of the mitochondrial pyruvate carrier Mpc3 was also significantly reduced in *pex3* cells (fold-change 0.66). We therefore wondered whether this could be related to dual localization on peroxisomes and mitochondria. This option was also inspired by recent reports on the dual localization of two other mitochondrial inner membrane carriers on peroxisomes and mitochondria *S. cerevisiae* Pet9, alias Aac2 (van Roermund et al., 2022) and *H. polymorpha* Mir1 (Pedersen et al., 2023 *preprint*). Moreover, although a peroxisomal pyruvate transporter was predicted to exist, because of the presence of the pyruvate producing enzyme Str3 in the matrix, it was never identified. Indeed, we established the dual localization of the heterodimeric Mpc1/Mpc3 pyruvate carrier to peroxisomes and mitochondria. This suggests that multiple mitochondrial carriers may be dually localized. Only a small fraction of Mpc1/Mpc3 was observed at a subset of the peroxisomes, which likely explains why the peroxisomal localization was missed in other studies, like high throughput FM (Yifrach et al., 2022) or organelle proteomics (Marelli et al., 2004). The presence of peroxisomal proteins in subdomains or on a subset of the peroxisomal population has also been reported before for other proteins (Cepińska et al., 2011; Yifrach et al., 2022).

In *S. cerevisiae*, expression of β-oxidation enzymes and Cta1 are known to be induced under peroxisome proliferation conditions. This effect is most pronounced for PTS1 and PTS2 enzymes whose expression is strongly inducible when peroxisome functions are needed (Karpichev and Small, 1998). In acetate grown yeast cells, genes encoding β-oxidation enzymes are derepressed (Kayikci and Nielsen, 2015). Notably, all 9 peroxisomal transcripts that were considerably enhanced in *pex3* cells encode peroxisomal β-oxidation enzymes and most of them were also increased at the protein level (**Figure 7**). In *pex3* cells, the β-oxidation pathway is most likely much less active due to the cytosolic mislocalization of the enzymes together with the lower abundance of Eci1 and Pox1, which produce the first product of the β-oxidation pathway (Dmochowska et al., 1990; Gurvitz et al., 1998). This will reduce the flux through the entire cascade of β-oxidation reactions, which explains why yeast *pex* mutants are unable to grow on oleic acid. As a result of the reduction in β-oxidation, the levels of free fatty acids may increase, leading to further induction of the expression of the genes encoding β-oxidation enzymes. Indeed, we observed that the transcripts of the β-oxidation enzymes were enhanced in *pex3* cells. The upregulation of β-oxidation enzymes might also be linked to the increased abundance of the receptor proteins Pex5 and Pex7 and/or may be a direct cellular response to counteract the loss of peroxisomes. Pox1 produces hydrogen peroxide. Hence, mislocalization of active Pox1 in the cytosol may cause oxidative stress. The considerably reduced levels of Pox1 protein in *pex3* cells may explain why we did not observe changes in transcripts/proteins of the cellular oxidative stress response. The reduced levels of Pox1 may be related to regulation by Arf1, which specifically controls protein but not transcript levels of Pox1 and Pxa1 post-translationally (Enkler et al., 2023). Unfortunately, we did not detect Pxa1 at the protein level. Possibly, Eci1 protein levels are controlled by the same mechanism. Our data further show that mislocalization of strongly upregulated β-oxidation enzymes in *pex3* cells resulted in enhanced levels of Hsp12 and Hsp31, which both play a role in early responses to prevent protein misfolding. This might also explain why levels of the molecular chaperone Hsp26, the disaggregase Hsp104, and the sequestrase Hsp42 were transcriptionally reduced in *pex3* cells.

The absence of peroxisomes not only resulted in changes in peroxisome related transcripts or proteins. Importantly, levels of several mitochondrial proteins were enhanced (**Figure 7**). These included proteins involved in mitochondrial respiration, protein import and protein refolding in the matrix. This may shed new light on the functional interactions between both cell organelles and may indicate that specific processes for safeguarding the fitness of mitochondria are induced in *pex3* cells. Moreover, it may give new hints to better understand the molecular mechanisms behind mitochondrial changes in peroxisome deficient human cells (Baumgart et al., 2001; Goldfischer et al., 1973; Nuebel et al., 2021).

Many other non-peroxisomal proteins were increased in *pex3* cells. An example is Tpo4 (a transporter of the major facilitator superfamily). Our further analysis did not provide any clues as to why this PM/vacuole transporter is enhanced in the absence of Pex3. Understanding why this protein is upregulated awaits further analysis. Also, genes that were strongly enhanced are very attractive for further analysis. The transcript with the highest increase in *pex3* cells is *EEB1*. This gene encodes acyl-coenzymeA:ethanol O-acyltransferase, which is implicated in lipid metabolism and toxification.

A large variety of poorly characterized proteins were reduced both at the transcript and protein level (e.g. Gre1, Sps100 Ykl107w). In addition, transcripts/proteins related to glucose metabolism, e.g. Tdh1 and Tdh2 (glyceraldehyde-3-phosphate dehydrogenases), Gnd2 6-phosphogluconate dehydrogenase) and Tkl2 (transketolase) were downregulated as well as Pma2 (a plasma membrane ATPase).

Interestingly, Gnd2 is a dually localized enzyme in the cytosol and peroxisomes in rat liver (Antonenkov, 1989). Furthermore, the role of Ykl107w, a member of the classical short-chain dehydrogenase/reductase family, in peroxisomal β-oxidation is conceivable. Thus, both enzymes are promising candidates for studying their potential function in peroxisome biology.

A few changes were also observed in proteins involved in the metabolism of non-preferred nitrogen sources. This includes enhanced protein levels of two transcription factors (Uga3 and Gzf3) that regulate genes involved in the uptake of these nitrogen sources. The amidolyase Dur12 is also regulated by Gzf3, however, the abundance of this protein was reduced in *pex3* cells. These data imply that in the absence of peroxisomes nitrogen metabolism has changed. This is intriguing because so far enzymes involved in nitrogen metabolism were never reported to exist in *S. cerevisiae* peroxisomes. This is in contrast to other yeast species, which contain several peroxisomal enzymes involved in the oxidation of organic nitrogen sources (e.g. amine oxidase, D-amino acid oxidase and urate oxidase in *H. polymorpha* (Veenhuis, 1992). Moreover, such enzymes also occur in peroxisomes of higher eukaryotes including plants (Tajima, 2004) and mammals (Angermüller et al., 2009).

We here show that the comparison of the whole cell proteome and transcriptome of a yeast *pex* mutant is a powerful approach to identifying novel processes and pathways influenced by peroxisomes. It would be interesting to also study other *pex* mutants that are not defective in PMP sorting, but in other processes. Attractive candidates include a *pex5 pex7* double mutant, which is selectively blocked in matrix protein import, or mutants that show defects in other aspects of peroxisome biology such as peroxisome fission (Pex11), peroxisome-ER contact site formation (Pex30), peroxisome inheritance (Inp1) or pexophagy (Atg36). Another option is to compare different *pex* mutants. Comparison of *pex3* and *pex19* cells may uncover new Pex3 or Pex19 functions. We already know that Pex3 is a multitasking protein that is also important for pexophagy by recruiting Atg36 and for peroxisome inheritance by recruiting Inp1 (Motley et al., 2012; Munck et al., 2009). In mammals, Pex19 is implicated in alternative functions unrelated to peroxisomes (Lyschik et al., 2022; Schrul and Kopito, 2016, Zimmermann et al., 2021). A comparison of *pex3* and *pex19* cells may therefore give novel hints to additional functions of these important peroxins in yeast.

Summarizing, we here show the power of the comparison of the whole cell proteome and transcriptome of a WT and peroxisome-deficient yeast mutant. The data obtained resulted in the identification of Mpc1/Mpc3 as another example of a dually localized mitochondrial carrier protein. Moreover, we uncovered many proteins/transcripts and cellular processes that have so far not been connected with peroxisome biology.

## Materials and methods

### Strains and growth conditions

The yeast strains used in this study are listed in **Table S3**. Yeast cells were grown in batch cultures on a mineral medium (MM) (Van Dijken et al., 1976) supplemented with 0.5 % glucose or 0.5 % acetate. When required, amino acids were added to the media at the following concentrations: 20 mg/L histidine, 30 mg/L leucine, 20 mg/L methionine, and 30 mg/L uracil. *S. cerevisiae* was grown at 30 °C, *H. polymorpha* at 37 °C.

For the selection of transformants, plates were prepared with 2 % agar in YPD (1 % yeast extract, 1 % peptone, and 1 % glucose) supplemented with 100 μg/mL zeocin (Invitrogen), 300 μg/mL hygromycin B (Invitrogen) or 100 μg/mL nourseothricin (Werner Bioagents).

*S. cerevisiae* cells were grown overnight in MM containing glucose, shifted to MM containing glucose, and shifted two times to MM containing acetate. For the analysis of (i) growth on acetate, the optical density of the cultures was measured at OD_660_ every 4 h with starting OD_660_ =0.1, (ii) acetate consumption, media was harvested at time points 0 h, 8 h, 12 h and 21 h, corresponding to OD_660_ of approximately 0.1, 0.5, 1.0 and 2.3 respectively; (iii) whole cell proteome and transcriptome, cells were harvested when cultures reached an OD_660_ of 0.5. For FM analysis *S. cerevisiae* cells were grown overnight in MM containing glucose, shifted to MM containing acetate twice, and imaged when reaching OD_660_ = 0.8-1.0. For spot assay, S. cerevisiae cells were grown overnight in MM containing glucose and shifted to MM containing acetate twice. When cells reached the OD_660_ = 1, serial dilutions were made (10^-1^, 10^-2^, 10^-3^, 10^-4^, 10^-5^) and spotted on mineral medium plates containing acetate (0.5%). Plates were incubated at 30 °C.

*H. polymorpha* cells were grown overnight in MM containing glucose, shifted to MM containing glucose twice, and imaged when reaching OD_660_ = 0.8-1.0.

### Measurement of acetic acid concentrations

Acetic acid was analyzed by HPLC equipped with a refractive index detector (Shimadzu, Kyoto, Japan) using an Aminex HPX-87H column (Bio-Rad) at a temperature of 65 °C using a mobile phase of 0.005 N H_2_SO_4_ and a flow rate of 0.55 ml/min.

### Molecular techniques

The primers, plasmids and DNA fragments used in this study are listed in **Tables S4**, **S5** and **S6**. DNA restriction enzymes were used as recommended by the suppliers (Thermo Fisher Scientific or New England Biolabs). Polymerase chain reactions (PCR) were carried out using Phusion High-Fidelity DNA polymerase or DreamTaq DNA polymerase (Thermo Fisher Scientific). DNA fragments were ligated using T4 DNA ligase (Thermo Fisher Scientific). For DNA and amino acid sequence analysis, Clone Manager 9 and SnapGene were used.

### Construction of *S. cerevisiae* strains

*S. cerevisiae* strains retrieved from different yeast libraries (Euroscarf) (Yofe et al., 2016; Weill et al., 2018; Meurer et al., 2018; Huh et al., 2003) were confirmed with PCR. To construct strains producing Pex14 mCherry, a PCR fragment encoding Pex14-mCherry was obtained using primers TER214 and TER215. Upon transformation, correct integration was confirmed by colony PCR using TER216 and R-mCherry-CP.

To obtain the *TPO4* deletion strains, a deletion cassette was amplified with primers F-Tpo4-del and R-Tpo4-del using pHIPH4 as a template. Replacement of the genomic *TPO4* gene in the *S. cerevisiae* strains with deletion cassette was confirmed with the colony PCR, using primers F-Tpo4-del-CP and R-Tpo4-del-CP, and Southern blotting.

Plasmid pHIPZ Tpo4 3xHA was obtained from GenScript, and constructed as follows: C-terminal fragment Tpo4 with 3xHA tag was synthesized and inserted into the pHIPZ mGFP-fusinator between *Hind*III and *Sph*I restriction sites. *Sfu*I restriction site was introduced to the fragment Tpo4 3xHA with silent mutation 1893C>G. *Sfu*I linearized pHIPZ Tpo4 3xHA was integrated into the TPO4 gene of *S. cerevisiae* WT and *pex3* deletion strain. Gene integration was confirmed with colony PCR using primers F_Tpo4 3xHA and pHIPZ-pHIPZ5 rew seq.

The PCR-based cloning approach and transformation of *S. cerevisiae* strains using the lithium acetate method were performed as previously described (Knop et al., 1999).

### Construction of *H. polymorpha* strains

To find homologs of *Sc* Mpc1 and Mpc3, we performed the blastp (NCBI) using all non-redundant protein sequence (nr) database in *H. polymorpha* (taxid:870730) and blastN, standard database in *Ogataea* (taxid:461281) (date used 6.3.2023). Two genes/proteins with the accession KAG7879638 and XP_013933851 were found, called here *Hp*Mpc1 and *Hp*Mpc3.

Plasmid pHIPZ18 Mpc1-meGFP was obtained as follows: PCR was performed on *H. polymorpha* WT genomic DNA using primers MPC1-C-fw and MPC1-C-rev. The PCR product was digested with *Hind*III and *Bam*HI, and inserted between the *Hind*III and *Bgl*II sites of the pHIPZ18 Inp1-meGFP. *Eco*RI linearized pHIPZ18 Mpc1-meGFP was integrated into the *ADH1* promoter of *H. polymorpha yku80* producing Pex3 mKate2, resulting in strain *Hp* P*_PEX3_* Pex3-mKate2::P*_ADH1_* Mpc1-meGFP. Gene integration was confirmed with colony PCR using primers cPCR MPC1-Cfw and mGFP rev check.

Plasmid pHIPZ18 Mpc3-meGFP was obtained as follows: PCR was performed on *H. polymorpha* WT genomic DNA using primers MPC3-C-fw and MPC3-C-rev. The PCR product was digested with *Hind*III and *Bgl*II, and inserted between the *Hind*III and *Bgl*II sites of the pHIPZ18 Inp1-meGFP. *Eco*RI linearized pHIPZ18 Mpc3-meGFP was integrated into the *ADH1* promoter of *H. polymorpha yku80* producing Pex3 mKate2, resulting in a strain *Hp* P*_PEX3_* Pex3-mKate2::P_ADH1_ Mpc3-meGFP. Gene integration was confirmed with colony PCR using primers cPCR MPC3-Cfw and mGFP rev check.

Transformations and site-specific integrations were performed as described previously (Faber et al., 1994).

### Cell lysis for proteomics analysis and proteolytic digestion

Cells were harvested by flash cooling the culture flasks on wet ice for 5 min followed by centrifugation for 7 min at 6000 g and 4°C. After harvesting, *S. cerevisiae* cells were washed once with deionized H_2_O, resuspended in 500 μl of urea buffer (8 M urea, 75 mM NaCl, 50 mM Tris-HCl, pH 8.0), and lysed by bead beating using a Minilys homogenizer (three cycles of 2 min at 4,000 rpm with 2 min cooling on ice between the cycles). For removal of cell debris, lysates were centrifuged for 5 min at 15,000 g and 4°C. Supernatants were collected and protein concentrations were determined using the bicinchoninic acid assay (Smith et al., 1985). 500 µg of protein per sample were treated with tris (2-carboxyethyl)phosphine (5 mM final concentration; 30 min at 37°C) to reduce cysteine residues followed by alkylation of free thiol groups using iodoacetamide (50 mM; 30 min at room temperature in the dark). Proteins were subsequently digested in solution using LysC and trypsin as previously described (Dannenmaier et al., 2018). Peptides were desalted using solid-phase extraction microspin disk cartridges (Affinisep, Le Houlme, France) that had been conditioned with methanol followed by 80% (v/v) acetonitrile (ACN)/0.5% (v/v) acetic acid and 0.5% (v/v) acetic acid. Peptides were loaded onto the cartridges, washed twice with 0.5% (v/v) acetic acid and eluted using 80% (v/v) ACN/0.5% (v/v) acetic acid. Solvents were evaporated and dried peptides were stored at -20°C until further use.

### Peptide stable isotope dimethyl labeling

Peptides (300 µg per sample) derived from WT and *pex3* cells were differentially labeled with ‘light’ formaldehyde (CH_2_O) and sodium cyanoborohydride (NaBH_3_CN) or the corresponding deuterated ‘heavy’ reagents (CD_2_O and NaBD_3_CN) as described before (Boersema et al., 2009; Peikert et al., 2017). Experiments were performed in three independent biological replicates. In replicate 1, peptides of WT and *pex3* cells were labeled ‘light’ and ‘heavy’, respectively. In replicates 2 and 3, the labeling was reversed to account for potential labeling bias. The labeling efficiency, assessed by LC-MS analysis, was > 97%. Equal amounts of differentially dimethyl-labeled peptides of WT and *pex3* cells (100 µg each) were mixed and dried *in vacuo*.

### High pH reversed-phase fractionation

To achieve high proteome coverage, peptide mixtures were fractionated using high pH reversed-phase liquid chromatography (LC) (Delmotte et al., 2007) as described before (Morgenstern et al., 2021). In brief, dried peptides were reconstituted in 100 µl of 99% solvent A (10 mM ammonium hydroxide, pH 10) and 1% solvent B (10 mM ammonium hydroxide in 90% ACN, pH 10), insoluble material was removed by centrifugation (12,000 x g, 5 min, room temperature), supernatants were filtered through a 0.2 mm PTFE membrane syringe filter (Phenomenex, Torrence, USA), and peptide mixtures were separated on an NX 3u Gemini C18 column (150 mm x 2 mm, particle size 3 µM, pore size 110Å; Phenomenex) operated at 40°C with a flow rate of 200 µl/min using an Ultimate 3000 HPLC system. Peptides were loaded at 1% solvent B (2 min) and eluted with a gradient of 1 – 50% B in 10 min, 50 – 78% B in 2 min and 1 min at 78% B. Fractions were collected in 50-s intervals (starting after 1 min, ending at 21 min) in a concatenated manner such that every 9th fraction was combined, which resulted in a total of 8 fractions per replicate. Peptides were dried *in vacuo*, desalted using StageTips as described before (Peikert et al., 2017), and acidified by adding trifluoroacetic acid to a final concentration of 0.1% (v/v).

### LC-MS analysis

Peptides were analyzed by nano-HPLC-tandem mass spectrometry (MS/MS) using an UltiMate 3000 RSLCnano system (Thermo Fisher Scientific, Dreieich, Germany) connected to an Orbitrap Elite mass spectrometer (Thermo Fisher Scientific, Bremen, Germany). The RSLC system was equipped with C18 pre-columns (nanoEase M/Z Symmetry C18; 20 mm length, 0.18 mm inner diameter; Waters) for washing and preconcentration of the peptides at a flow rate of 10 µl/min and an analytical C18 reversed-phase nano LC column (nanoEase M/Z HSS C18 T3; 250 mm length, 75 mm inner diameter, 1.8 µm particle size, 100 Å packing density; Waters) for peptide separation (flowrate, of 300 nl/min). Peptides were eluted using a binary solvent system that consisted of 4% (v/v) dimethyl sulfoxide (DMSO)/0.1% (v/v) formic acid (FA) (solvent A) and 4% (v/v) DMSO/0.1% (v/v) FA/30% (v/v) ACN/48% (v/v) methanol (solvent B). Peptides equivalent to 1 µg of protein were loaded onto the pre-columns for 5 min using solvent A and eluted applying a gradient ranging from 1 – 7% solvent B in 5 min, 7 – 52% B in 145 min, 52% – 95% B in 85 min, and 5 min at 95% B.

Mass spectrometric data were acquired in data-dependent mode using the following parameters: mass range of *m/z* 370 – 1700 for MS precursor scans at a resolution of 120,000 (at *m/z* 400); maximum automatic gain control (AGC) of 10^6^ ions; maximum injection time (IT) of 200 ms. A TOP20 method was applied for low-energy collision-induced dissociation of precursor ions with a normalized collision energy of 35%, an activation q of 0.25, and an activation time of 10 ms. The maximum AGC for MS/MS scans was set to 5 × 10^3^ ions, the maximum IT to 150 ms, and the dynamic exclusion time to 45 sec.

### MS data analysis

For protein identification and relative quantification, the MaxQuant software package (version 1.6.10.43) (Cox and Mann, 2008) with its integrated search engine Andromeda (Cox et al., 2011) was employed. MS/MS data were searched against the *S. cerevisiae* reference proteome provided by UniProt (www.uniprot.org/proteomes/UP000002311; downloaded April 2021; 6,079 entries). The database search was carried out using trypsin and LysC as proteolytic enzymes, allowing a maximum of three missed cleavages, and applying mass tolerances of 4.5 ppm for precursor and 0.5 Da for fragment ions. Carbamidomethylation of cysteine residues was set as fixed modification, oxidation of methionine and acetylation of protein N-termini were considered as variable modifications, and DimethLys0/DimethNter0 and DimethLys6/DimethNter6 were selected as light and heavy labels, respectively. The options ’match between runs’ and ‘requantify’ were enabled. ‘Requantify‘ enables the calculation of protein abundance ratios in experiments that employ stable isotope labeling if only the labeled or unlabeled variant of a peptide is present. For example, this is the case for Pex3 peptides, which are absent in *pex3* cells. Using the ‘requantify’ option, the algorithm assigns a peptide intensity for the missing counterpart from the background signals in MS spectra at the expected *m/z* value. Protein identification was based on ≥ one unique peptide with a minimum length of seven amino acids. A false discovery rate of 0.01 was applied at both peptide and protein levels. For relative protein quantification, ≥ one ratio count (considering unique and razor peptides) was required.

For data analysis and visualization, MaxQuant results were processed using the Python-based analysis pipeline "autoprot" (Bender et al., 2024 p*reprint*). Only proteins quantified in at least two out of three replicates were considered for further analysis. Normalized protein abundance ratios calculated by MaxQuant were log_2_-transformed, followed by sequential imputation of missing values and cyclic-loess normalization (Cleveland and Devlin, 1988). To identify proteins with differences in protein abundance between WT and *pex3* cells, we used the ‘linear models for microarray data’ (limma) approach, a moderated two-sided t-test that adjusts a protein’s variance in ratios between replicates towards the average ratio variance of the entire dataset (Smyth, 2004; Schwämmle et al., 2013). Information about proteins identified and quantified are provided in **Tables S1a** and **S1c**. Information about peroxisomal localization of proteins or association with peroxisomes, used to assess the effect of *PEX*3 deletion on peroxisomal proteins and transcripts, is based on published data (David et al., 2022; Elbaz-Alon et al., 2014; Marelli et al., 2004; Motley et al., 2012; Tanaka et al., 2014; Yifrach et al., 2022).

### Transcriptomics analysis

Cells were harvested by flash cooling the culture flasks on wet ice for 5 min followed by centrifugation for 7 min at 6000 g at 4°C. The supernatants were decanted, and pellets were subjected to an additional 1 min centrifugation to remove residual supernatant. After removal of the supernatant, the pellets were solubilized in RNA*later*® (Sigma, #R0901) according to the instructions of the manufacturer and incubated on wet ice for 5 min. The samples were spun down for 10 min at 6000 g at 4°C and the supernatants were removed. The samples were centrifuged for an additional 1 min at 6000 g to remove residual supernatant and the cell pellets were flash frozen in liquid nitrogen, stored at -80 °C and shipped on dry ice for further analysis.

Transcriptomic analysis was performed by Eurofins as a service package named NGSelect RNA analysis (Eurofins Germany, #6WB2-RNAI01). 150 ng total mRNA was used. The mRNA was poly-A purified and fragmented, followed by random primed cDNA library synthesis, adapter ligation and adapter specific PCR amplification. Sequencing was performed using the Illumina BIO-IT platform, as paired end with 5 million read pairs (2 x 150 bp) per sample. For reference, the *Saccharomyces cerevisiae* S288c assembly from the *Saccharomyces* Genome Database R64-1-1 (INSDC Assembly GCA_000146045.2, Sep 2011) was used (Liachko et al., 2013).

For data analysis, genes with less than 10 reads on average across the compared groups were removed. The abundance counts of each gene were then used to perform differential gene expression (DGE) analysis (Oshlack et al., 2010). DGE was performed using the R/Bioconductor DESeq2 package (Love et al., 2014). Standard normalization and differential expression analysis steps were subjected to a single function in DESeq2m using the function DESeqDataSetFromMatrix. P-values were calculated using the Wald test implemented in the R/Bioconductor DESeq2 package.

### Gene Ontology term enrichment analysis

GO term enrichment analyses of proteins and transcripts exhibiting a minimum fold-change of 1.5 (reduced/increased; p-value < 0.05) in *pex3* compared to WT cells were performed using the shinyGO application (version 0.77; http://bioinformatics.sdstate.edu/go/)(Ge et al., 2020) and all proteins and transcripts identified in our omics analysis as background. False discovery rate p-values were corrected for multiple testing using the Benjamini-Hochberg method implemented in the shinyGO analysis pipeline. GO terms with corrected p-values of < 0.05 were considered enriched. The results of the GO term enrichment analyses are provided in **Table S2**.

### Fluorescence microscopy

Widefield fluorescence microscopy images of living cells were captured in a growth medium at room temperature. Images were obtained using an Axioscope A1 microscope (Carl Zeiss), with a 100 × 1.30 NA objective, Micro-Manager 1.4 software, and a digital camera (Coolsnap HQ2; Photometrics). GFP and mNeonGreen fluorescence was visualized using a 470/40 nm band-pass excitation filter, a 495 nm dichromatic mirror, and a 525/50 nm band-pass emission filter. mKate2 and mCherry fluorescence was visualized using a 587/25 nm band-pass excitation filter, a 605 nm dichromatic mirror, and a 670/70 nm band-pass emission filter.

For Airyscan imaging, cells were fixed in 1% formaldehyde for 30 min on ice and incubated in 10 mM glycine in 100 mM PBS (pH 7.5) for 60 min on ice. Airyscan images were captured with a confocal microscope (LSM800; CarlZeiss) equipped with a 32 channel gallium arsenide phosphide photomultiplier tube (GaAsP-PMT), Zen 2009 software (CarlZeiss), and a 63×1.40 NA objective (CarlZeiss). The GFP signal was visualized by excitation with a 488 nm laser, and mCherry was visualized with a 561 nm laser.

Image analysis was performed using ImageJ, and all brightfield images have been adjusted to only show cell outlines.

### Immunoblotting

Total cell extracts were prepared from trichloroacetic acid (TCA) treated cells as described previously (Baerends et al., 2000; McCammon et al., 1994). Sodium dodecyl sulfate–polyacrylamide gel electrophoresis (SDS-PAGE) was performed as described previously (Laemmli, 1970; Baerends et al., 2000). Equal volumes of samples were loaded per lane.

Semi-dry transfer of proteins from the polyacrylamide gel to a nitrocellulose membrane (Amersham Protran 0.2 NC, #10600001) was performed as described before (Kyhse-Andersen, 1984). Blots were decorated with anti-pyruvate carboxylase-1 (Pyc1) antibodies (1:10,000 dilution) (Ozimek et al., 2007), anti-Pot1 (1:10,000 dilution), anti-Pex14 (1:5,000 dilution) (Niederhoff et al., 2005), anti-Pex5 (1:10,000 dilution) (Albertini et al., 1997), or anti-HA (1:10,000 dilution) (Roche, #12CA5). Secondary goat anti-rabbit (#31460) or goat anti-mouse (#31430) antibodies conjugated to horseradish peroxidase (HRP) (Thermo Scientific; 1:5,000 dilution) were used for detection. Chemiluminescent (Amersham ECL Prime, #RPN2232) detection solution was performed according to the manufacturer’s guidelines (Bio-Rad, Chemidoc Imaging System). Blots were imaged and analyzed using the Bio-Rad ChemiDoc imaging System.

## Data Availability Statement

The mass spectrometry proteomics data have been deposited to the ProteomeXchange Consortium (Deutsch et al., 2023) via the PRIDE (Perez-Riverol et al., 2022) partner repository and are accessible using the dataset identifier PXD047234. The transcriptome data were deposited at NCBI-GEO (Project accession GSE261289).

## Supporting Information

This article contains supporting information.

## Acknowledgments

We thank Maya Schuldiner and Einat Zalckvar (Weizmann Institute) for making yeast strains available and Ralf Erdmann (Ruhr University Bochum) for antibodies. We are grateful to Jeroen Nijland for performing the acetate measurements and to Rinse de Boer for assistance in preparing the microscopy images. We thank Arjan Krikken and Jan Kiel (RUG) for helpful discussions and expert technical assistance. This project received funding from the European Union’s Horizon 2020 research and innovation programme under the Marie Skłodowska-Curie grant agreement No 812968.

## Author contributions

IvdK and BW conceived the project and supervised the study. TK, MPP, HD and MA performed the experiments. TK and MPP grew and harvested the cells. HD, SO performed proteomics and quantitative MS data analysis. Analysis of the transcriptomics data was performed by MA and HD. HD, SO and MA performed bioinformatics and statistical analysis. TK performed the acetate measurements and yeast growth analysis. TK constructed yeast strains, performed fluorescence microscopy and spot assays. MPP, TK made western blots. TK, IvdK and BW wrote the original draft of the manuscript. TK, MPP, HD and SO prepared figures and tables. All authors contributed to reviewing and editing the manuscript. All authors approved the final version of the manuscript.

