## Supporting figures and tables for "Integrative Omics reveals changes in the cellular landscape of yeast without peroxisomes"

<u>Material included:</u>

Figures S1 - S3

Supporting Table Legends (Tables S1 and S2; xlsx files)

Supporting Tables S3 - S6

References

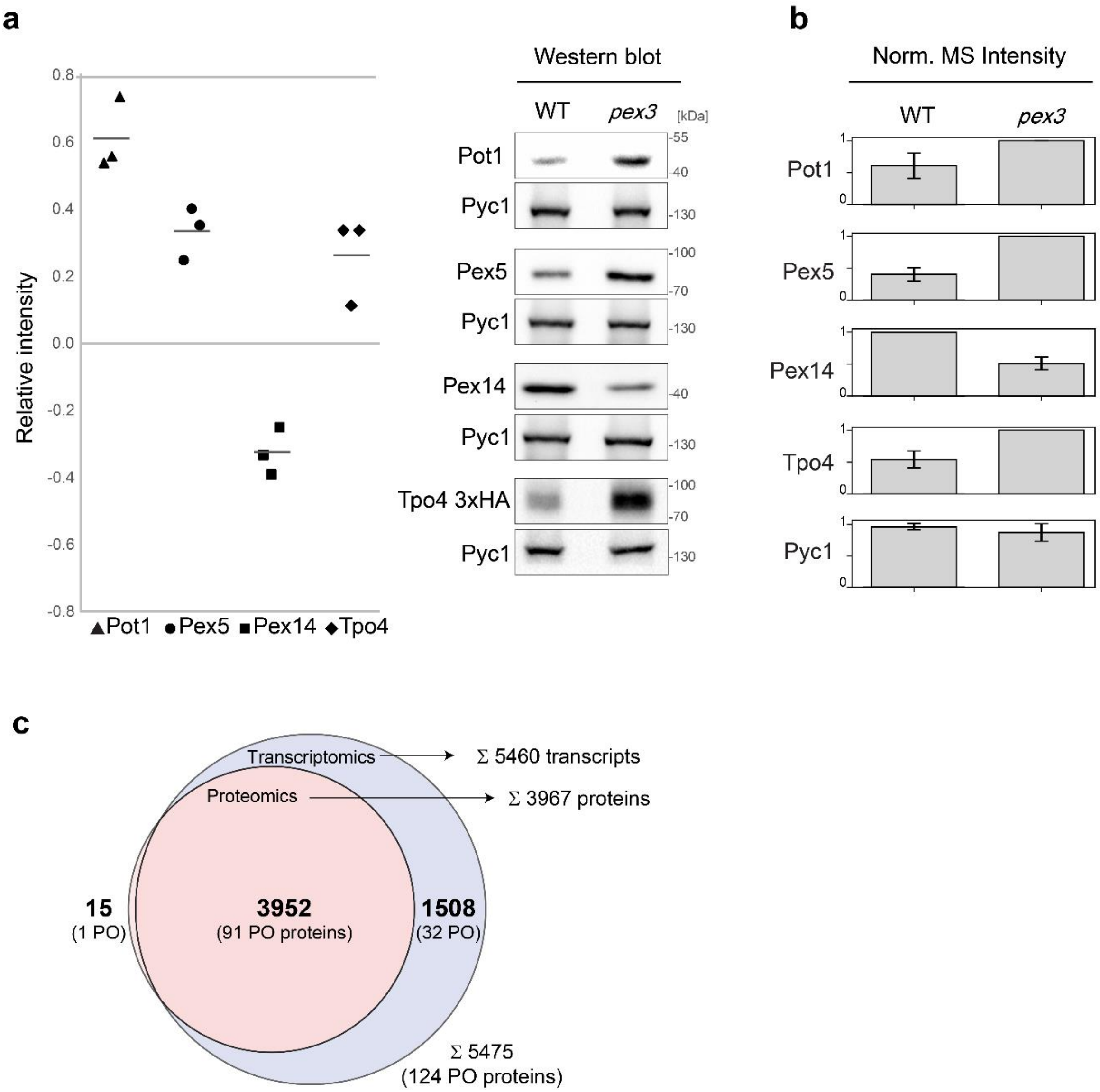


**Figure S1. Validation of selected regulated proteins and comparison of proteomics *versus* transcriptomics data.**

(a) Validation of differences in protein abundance in whole cell lysates of wild-type (WT) and *pex3* cells by Western blotting using antibodies specifically recognizing the indicated proteins. HA-tagged Tpo4 was recognized using an anti-HA antibody. Pyruvate carboxylase 1 (Pyc1) served as loading control. Signal intensities of three independent replicates were quantified by densitometry. Relative protein abundances for WT and *pex3* cells were determined by normalizing the signal intensities of Pot1, Pex5, Pex14, and Tpo4 to the signal of the corresponding Pyc1 control. The scatter plot shows the log-transformed ratios between the relative protein abundances determined for *pex3* and WT cells (*i.e.*, relative intensity) for the different proteins.

(b) Normalized MS intensities for the proteins selected for Western blot validation in (a). Data are extracted from the proteomics analysis of WT *versus pex3* cells (Table S1a).

(c) Overlap of proteins and transcripts identified in this study in the proteomics and transcriptomics analysis. Please note that the number of individual proteins reported in this figure (*i.e.*, 3967 in total) is higher than the number of protein groups (3921) that were identified in the proteomics analysis. This is due to the matching of the transcriptomics data to the proteomics data. Several of the protein groups reported by the analysis software MaxQuant contain more than one protein (see Table S1a) because the identified peptides do not allow to discriminate between the proteins within the protein group. However, we obtained data for several transcripts of distinct proteins that were combined in a protein group. Hence, these proteins with matching transcripts are considered and counted here as individual proteins (see also Tables S1c and S1d). PO, peroxisomal.

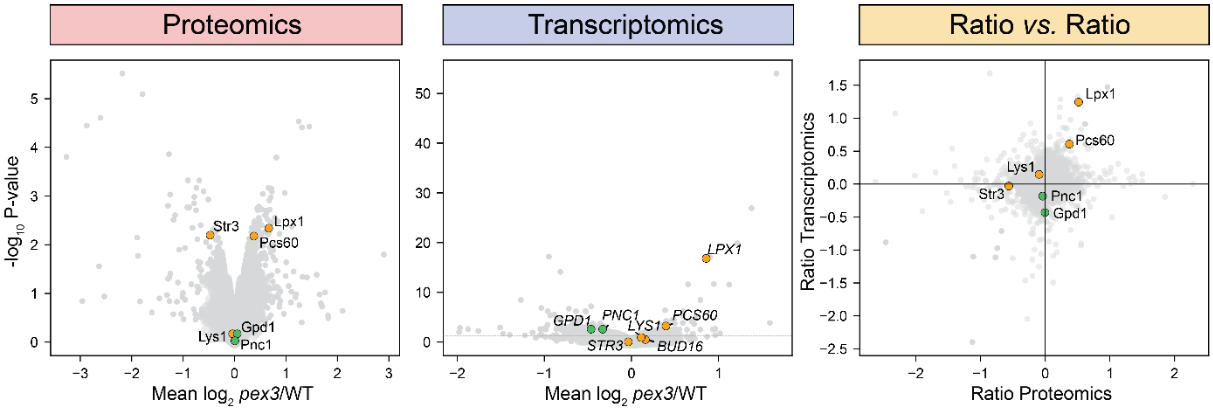


**Figure S2. Effect of Pex3 deficiency on peroxisomal enzymes and their transcripts associated with diverse metabolic processes.** Same plots as shown in Figure 3**.** Stress-inducible peroxisomal enzymes are marked in green.

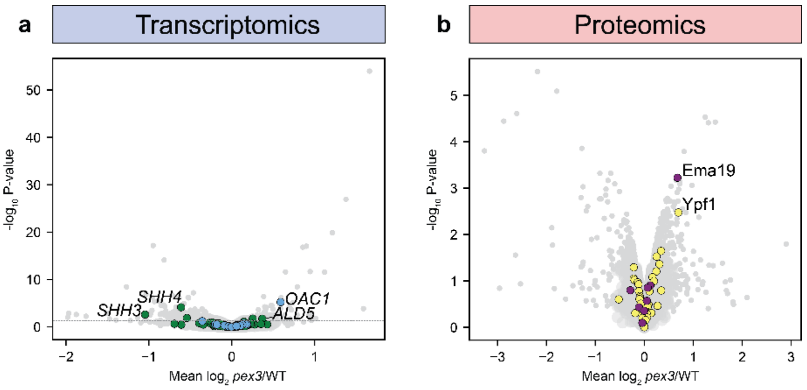


**Figure S3. Abundance of mitochondrial transcripts (a), ERAD and ER-SURF components (b) in *pex3 versus* WT cells.** (a) Same plot as shown in Figure 2c highlighting transcripts of mitochondrial respiratory chain components (green), mitochondrial carrier, and mitochondrial import proteins (both in blue). (b) Same plot as shown in Figure 2b highlighting proteins associated with the endoplasmic reticulum unfolded protein response (ERAD) (yellow) and ER-SURF (purple) (Koch et al., 2021). A list of ERAD components was retrieved from the *Saccharomyces* Genome Database (https://www.yeastgenome.org/) using the Gene Ontology identifier GO:0036503. Transcripts/proteins annotated in the subplots are discussed in the manuscript.

### Supporting Tables

**Table S1. Results of quantitative proteomics and transcriptomics analyses of *Saccharomyces cerevisiae* wild-type *versus pex3* cells** (xlsx file)

(a) Whole cell extracts of *Saccharomyces cerevisiae* wild-type (WT) and *pex3* cells were analyzed by quantitative mass spectrometry using peptide stable isotope dimethyl labeling (n=3). Raw MS data and complete MaxQuant results files are available via Proteome-Xchange with the identifier PXD047234.

(b) Transcriptomics data obtained from mRNA extracted from the same samples that were used for the proteomics analysis.

(c) Combined omics data.

(d) Heat map of proteomics and transcriptomics data visualizing *pex3*/WT ratios ($log_2$ fold-changes) for all proteins/transcripts identified in this study.

**Table S2. Results of GO term enrichment analysis** (xlsx file)

GO term enrichment analysis for the domains "Biological Process" and "Cellular Component" was performed for proteins/transcripts with a minimum fold-change of 1.5 (reduced and increased; p-value < 0.05) between wild-type and *pex3* cells using shinyGO (version 0.77; http://bioinformatics.sdstate.edu/go/). GO terms with Benjamini-Hochberg corrected p-values of < 0.05 were considered enriched.

**Table S3. Yeast strains used in this study.**

| Strain | Description and genotype | Reference |
|---|---|---|
| *Sc* BY4741 (WT) | MATa his3Δ1 leu2Δ0 met15Δ0 ura3Δ0 | Euroscarf #Y00000 |
| *Sc* BY4741 *pex3* | MATa his3Δ1 leu2Δ0 met15Δ0 ura3Δ0 pex3(YDR329c)::kanMX4 | Euroscarf #Y03688 |
| *Sc* WT $P_{NOP1}$ sfGFP-Mdh3 | MATa can1Δ::GAL1pr-SceI::STE2pr-SpHIS5 his3Δ1 leu2Δ0 met15Δ0 ura3Δ0 hphΔn::URA3::SpNOP1pr-sfGFP-Mdh3 | (Weill et al., 2018; Yofe et al., 2016 |
| *Sc* WT $P_{NOP1}$ sfGFP-Cat2 | MATa can1Δ::GAL1pr-SceI::STE2pr-SpHIS5 his3Δ1 leu2Δ0 met15Δ0 ura3Δ0 hphΔn::URA3::SpNOP1pr-sfGFP-Cat2 | (Weill et al., 2018; Yofe et al., 2016) |
| *Sc* WT $P_{NOP1}$ sfGFP-Cit2 | MATa can1Δ::GAL1pr-SceI::STE2pr-SpHIS5 his3Δ1 leu2Δ0 met15Δ0 ura3Δ0 hphΔn::URA3::SpNOP1pr-sfGFP-Cit2 | (Weill et al., 2018; Yofe et al., 2016) |
| *Sc* WT $P_{MDH3}$ Mdh3-mNG | MATa can1Δ::GAL1pr-SceI::STE2pr-SpHIS5 his3Δ1 leu2Δ0 met15Δ0 ura3Δ0 lys2+/lys+ lyp1Δ::STE3pr-LEU2 Mdh3-mNeonGreen-ADH1term:Hygro | (Meurer et al., 2018) |
| *Sc* WT $P_{CAT2}$ Cat2-mNG | MATa can1Δ::GAL1pr-SceI::STE2pr-SpHIS5 his3Δ1 leu2Δ0 met15Δ0 ura3Δ0 lys2+/lys+ lyp1Δ::STE3pr-LEU2 Mdh3-mNeonGreen-ADH1term:Hygro | (Meurer et al., 2018) |
| *Sc* WT $P_{CIT2}$ Cit2-GFP | MATa his3Δ1 leu2Δ0 met15Δ0 ura3Δ0; Cit2-GFP HIS3MX6 | (Huh et al., 2003) |
| *Sc* WT $P_{MPC1}$ Mpc1-GFP | MATa his3Δ1 leu2Δ0 met15Δ0 ura3Δ0; Mpc1-GFP HIS3MX6 | (Huh et al., 2003) |
| *Sc* WT $P_{MPC1}$ Mpc1-GFP:: $P_{PEX14}$ Pex14-mCherry | *Sc* $P_{MPC1}$ Mpc1-GFP with integration of Pex14 mCherry fragment | This study |
| *Sc* WT $P_{MPC3}$ Mpc3-GFP | MATa his3Δ1 leu2Δ0 met15Δ0 ura3Δ0; Mpc3-GFP HIS3MX6 | (Huh et al., 2003) |
| *Sc* WT $P_{MPC3}$ Mpc3-GFP:: $P_{PEX14}$ Pex14-mCherry | *Sc* $P_{MPC3}$ Mpc3-GFP with integration of Pex14 mCherry fragment. | This study |
| *Sc* WT $P_{TPO4}$ Tpo4-GFP | MATa his3Δ1 leu2Δ0 met15Δ0 ura3Δ0; Tpo4-GFP HIS3MX6 | (Huh et al., 2003) |
| *Sc* WT $P_{TPO4}$ Tpo4-GFP:: $P_{PEX14}$ Pex14-mCherry | *Sc* $P_{TPO4}$ TPO4-GFP with integration of Pex14 mCherry fragment | This study |
| *Sc* WT $P_{TPO4}$ Tpo4 3xHA | *Sc* BY4741 (WT) with integration of plasmid pHIPZ Tpo4 3xHA. | This study |
| *Sc pex3* $P_{TPO4}$ Tpo4 3xHA | *Sc* BY4741 *pex3* with integration of plasmid pHIPZ Tpo4 3xHA. | This study |
| *Sc tpo4* | *Sc* BY4741 (WT) with integration of *tpo4* deletion cassette. | This study |

| | | |
|---|---|---|
| *Sc pex3 tpo4* | *Sc* BY4741 *pex3* with integration of *tpo4* deletion cassette. | This study |
| *Hp* NCYC495 (WT) | *yku80*::URA3; *leu1.1* | (Saraya et al., 2012) |
| *Hp* WT $P_{PEX3}$ Pex3-mKate2 | *Hp* WT with integration of plasmid pHIPN Pex3-mKate2. | (Krikken et al., 2020) |
| *Hp* WT $P_{PEX3}$ Pex3-mKate2:: $P_{ADH1}$ Mpc1-meGFP | *Hp* WT $P_{PEX3}$ Pex3-mKate2 with integration of plasmid pHIPZ18 Mpc1-meGFP. | This study |
| *Hp* WT $P_{PEX3}$ Pex3-mKate2:: $P_{ADH1}$ Mpc2-meGFP | *Hp* WT $P_{PEX3}$ Pex3-mKate2 with integration of plasmid pHIPZ18 Mpc2-meGFP. | This study |

**Table S4. Plasmids used in this study.**

| Plasmid | Description | Reference |
|---|---|---|
| pHIPH Pex14-mCherry | Plasmid containing C-terminal region of Sc *PEX14* fused to mCherry ($Hph^R$, $Amp^R$). | (Thomas et al., 2018) |
| pHIPN Pex3-mKate2 | Plasmid containing C-terminal region of *Hp PEX3* fused to mKate2 ($Nat^R$, $Amp^R$). | (Krikken et al., 2020) |
| pHIPZ18 Inp1-meGFP | Plasmid for integration of genes the *ADH1* promoter ($Zeo^R$, $Amp^R$). | (Krikken et al., 2020) |
| pHIPZ18 Mpc1-meGFP | Plasmid containing *Hp MPC1* fused to mEGFP under the *ADH1* promoter ($Zeo^R$, $Amp^R$). | This study |
| pHIPZ18 Mpc2-meGFP | Plasmid containing *Hp MPC2* fused to mEGFP under the *ADH1* promoter ($Zeo^R$, $Amp^R$). | This study |
| PHIPZ mGFP-fusinator | Plasmid used for the introduction of Tpo4 3xHA fragment ($Zeo^R$, $Amp^R$). | (Saraya et al., 2010) |
| PHIPZ Tpo4 3xHA | Plasmid containing C-terminal region of *Sc TPO4* fused to 3xHA ($Zeo^R$, $Amp^R$). | This study |

**Table S5. Primers used in this study.**

| Name | Sequence |
|---|---|
| R-MDH3-CP | CAACAGCGGTTACCAGGC |
| R-CAT2-CP | CCTTACATAACAGCTGCTGTCT |
| R-CIT2-CP | GGTTGTGAGCTTCCTTTTGC |
| F-sfGFP-CP | TCAGGCACAACGTCGAAGAT |
| F-MDH3-CP | GGCCAAGTTTGCTGAAGAAG |
| F-CAT2-CP | GGTAATGGTGTCGATCGTCA |
| R-mNG-CP | CCGATATGAGGGACCAGTATCC |
| F-CIT2-CP | TGCCATGGACCATTTTCCAG |
| R-GFP(S65T)-CP | CAACAAGAATTGGGACAACTCC |
| F_Prom-Pex3_Sc | GGTAGTTAATACTAGTCATCGT |
| R_Pex3_Sc | TGGTGTGAGTGTCAGTAC |
| R_KanMX4 | ATCATCAGGAGTACGGATA |
| F-Mpc1-CP | CAACCGGTTCAACGCG |
| F-Mpc3-CP | GGTCTTCGCAGGGCTAAATG |
| TER214 | AGATCAGTGTCCCTGACTGGCAAAATGGACAGGTCGAAG ACTCCATCCCAATGGTGAGCAAGGGCGAGGAGGAT |
| TER215 | GTTACAATTACAATTTCCGTTAAAAAACTAATTACTTACATA GAATTGCGCGTTTTCGACACTGGATGGCGGCGTT |
| TER216 | TAACCGTATGGAATCCGGTA |
| R-mCherry-CP | GGCCTTGGAGCCGTACATGA |
| MPC1-C-fw | CCCAAGCTTATGAGCTCGTCCTCGGCGTG |
| MPC1-C-rev | CGCGGATCCTTTCTGCACCTTGTTGCCGT |
| cPCR MPC1-Cfw | CACGGCAGCAGAATTGGAATTG |
| MPC3-C-fw | CCCAAGCTTATGGCCACGGCATTCAGAAG |
| MPC3-C-rev | GGAAGATCTTGCCTCTGTTTTCGCTGGTGTAGTGC |
| cPCR MPC3-Cfw | CACGGCAGCAGAATTGGAATTG |
| mGFP rev check | AAGTCGTGCTGCTTCATGTG |
| F_Tpo4 3xHA | CGTAAATGCTGTCGATGGTGAATTG |
| pHIPZ-pHIPZ5 rew seq. | CGAGGTTTTGTCCTTGGTCT |
| F-Tpo4-del-CP | GACACATCGATTTTCTAGTCATCAC |
| R-Tpo4-del-CP | GAATTTTATCCAGCTATCCCCC |

**Table S6. DNA fragment used in this study.**

| Name | Sequence |
|---|---|
| Tpo4 3xHA | GGTGAATTGAATCTAACAAGGATGACCACGTTAAGGACCATGGAAACAG<br>ACCCTTCGACTAGAGAAAAACCAGGTGAAAGGCTATCTCTGCGCAGGAC<br>CCATACGCAGCCCGTTCCTGCCTCGTTTGATCGCGAGGACGGGCAACATG<br>CGCAAAATCGTAATGAACCCATTTCGAATAGTTTATACTCAGCTATCAAGG<br>ATAATGAAGACGGTTATTCGTATACGGAAATGGCCACCGATGCTTCCGCC<br>AGAATGGTTTACCCATACGACGTTCCAGATTACGCTTACCCATACGACGT<br>TCCAGATTACGCTTACCCATACGACGTTCCAGATTACGCTTGA |
